# A computational knowledge engine for human neuroscience

**DOI:** 10.1101/701540

**Authors:** Elizabeth Beam, Christopher Potts, Russell A. Poldrack, Amit Etkin

## Abstract

Functional neuroimaging has been a mainstay of human neuroscience for the past 25 years. The goal for this research has largely been to understand how activity across brain structures relates to mental constructs and computations. However, interpretation of fMRI data has often occurred within knowledge frameworks crafted by experts, which have the potential to reify historical trends and amplify the subjective biases that limit the replicability of findings. ^1^ In other words, we lack a comprehensive data-driven ontology for structure-function mapping in the human brain, through which we can also test the explanatory value of current dominant conceptual frameworks. Ontologies in other fields are popular tools for automated data synthesis,^2, 3^ yet relatively few attempts have been made to engineer ontologies in a data-driven manner.^4^ Here, we employ a computational approach to derive a data-driven ontology for neurobiological domains that synthesizes the texts and data of nearly 20,000 human neuroimaging articles. The data-driven ontology includes 6 domains, each defined by a circuit of brain structures and its associated mental functions. Several of these domains are omitted from the leading framework in neuroscience, while others uncover novel combinations of mental functions related to common brain circuitry. Crucially, the structure-function links in each domain better replicate across articles in held-out data than those mapped from the dominant frameworks in neuroscience and psychiatry. We further show that the data-driven ontology partitions the literature into modular subfields, for which the domains serve as generalizable archetypes of the structure-function patterns observed in single articles. The approach to computational ontology we present here is the most comprehensive functional characterization of human brain circuits quantifiable with fMRI. Moreover, our methods can be extended to synthesize other scientific literatures, yielding ontologies that are built up from the data of the field.

---

Human neuroimaging seeks to understand how mental processes interrelate with patterns of brain activity. The flow of inquiry, however, has been largely unidirectional – taking mental constructs defined decades earlier in psychology as the premise for brain mapping efforts. As a consequence, neuroimaging studies have often served to reify theorized distinctions between psychological constructs, rather than to derive novel constructs anchored on brain function.^5^ Meta-analyses summarizing results across task paradigms have in many cases shown that brain circuits measured at the resolution of fMRI are surprisingly more similar than different between psychological constructs.^6, 7, 8^ In turn, constructs accepted as “natural kinds” have been mapped to heterogeneous neural activation patterns across the literature.^9, 10, 11^ Nonetheless, these meta-analyses have been limited to only addressing a subset of mental constructs or neural structures. By applying natural language processing (NLP) and machine learning techniques to the collective results of 25 years of human neuroimaging, we sought here to redefine mental constructs in relation to latent patterns in brain activation data, yielding an integrative data-driven ontology for neurobiological domains.

An empirical description of brain function is needed not only to guide the next generation of basic human neuroscience, but also as a biological foundation for classifying psychiatric disease. It is now well appreciated that the symptom-based mental illness diagnoses specified in the *Diagnostic and Statistical Manual* (DSM) are highly comorbid, biologically heterogeneous, and poorly predictive of treatment response.^12, 13^ The field of biological psychiatry has attempted to address these shortcomings by reframing mental illnesses as variations in the function of basic brain systems. This has yielded one of the most comprehensive and conceptually dominant expert-driven frameworks for mapping circuits, behaviors, and other units of analysis – called the Research Domain Criteria (RDoC) project.^13^ However, the organizational principles of RDoC remain largely untested, the reproducibility of its circuit-function links is unknown, and the performance of this expert-driven framework relative to a fully data-driven one has not been investigated. To assess the utility of RDoC and the DSM in explaining structure-function relationships in the human brain, and to contrast these frameworks with a data-driven ontology, we conducted a comprehensive neuroimaging meta-analysis of 18,155 PET and fMRI articles.

In engineering a data-driven ontology for human neuroscience, we sought to identify the most coherent set of mental functions (from article texts) for describing the most distinct functional brain circuits (from coordinate data), which together comprise what we refer to as neurobiological “domains” (Fig. 1a). The corpus of 18,155 articles used to generate the data-driven ontology was curated based on the availability of coordinate data from one of several sources, including the BrainMap^14^ and Neurosynth^15^ databases (Extended Data Fig. 1a). To define the set of possible mental functions, we compiled a broad and diverse lexicon of 1,683 words and phrases from public sources such as RDoC,^12^ the BrainMap Taxonomy,^14^ and the Cognitive Atlas^16^ (Extended Data Fig. 2). The lexicon includes terms for mental constructs (e.g., “emotional memory”), processes (e.g., “retrieval”), percepts and stimuli (e.g., “face”), and task paradigms (e.g., “face identification task”). The mental function terms were extracted from the preprocessed full texts of articles in our corpus, which contained 128,170,267 total words and 4,831,488 occurrences of terms from our lexicon. We next mined the neuroimaging article data, which included 605,292 spatial coordinates in the human brain. Coordinates were mapped probabilistically in a one-to-many fashion onto 114 gray matter structures in a neuroanatomical atlas spanning the human cerebrum^17^ and cerebellum,^18^ resulting in 2,636,325 total labels for brain structures. Data were randomized and split into a training set containing 70% of articles (*n* = 12,708) for generating the data-driven ontology and fitting models, a validation set containing 20% of articles (*n* = 3,631) for optimizing model hyperparameters and selecting thresholds for the data-driven ontology, and a test set containing 10% of articles (*n* = 1,816) for comparing the data-driven ontology against RDoC and the DSM. Splitting the data *a priori* enabled us to avoid biased estimation of the reproducibility of circuit-function links in subsequent analyses.

**Fig. 1.**
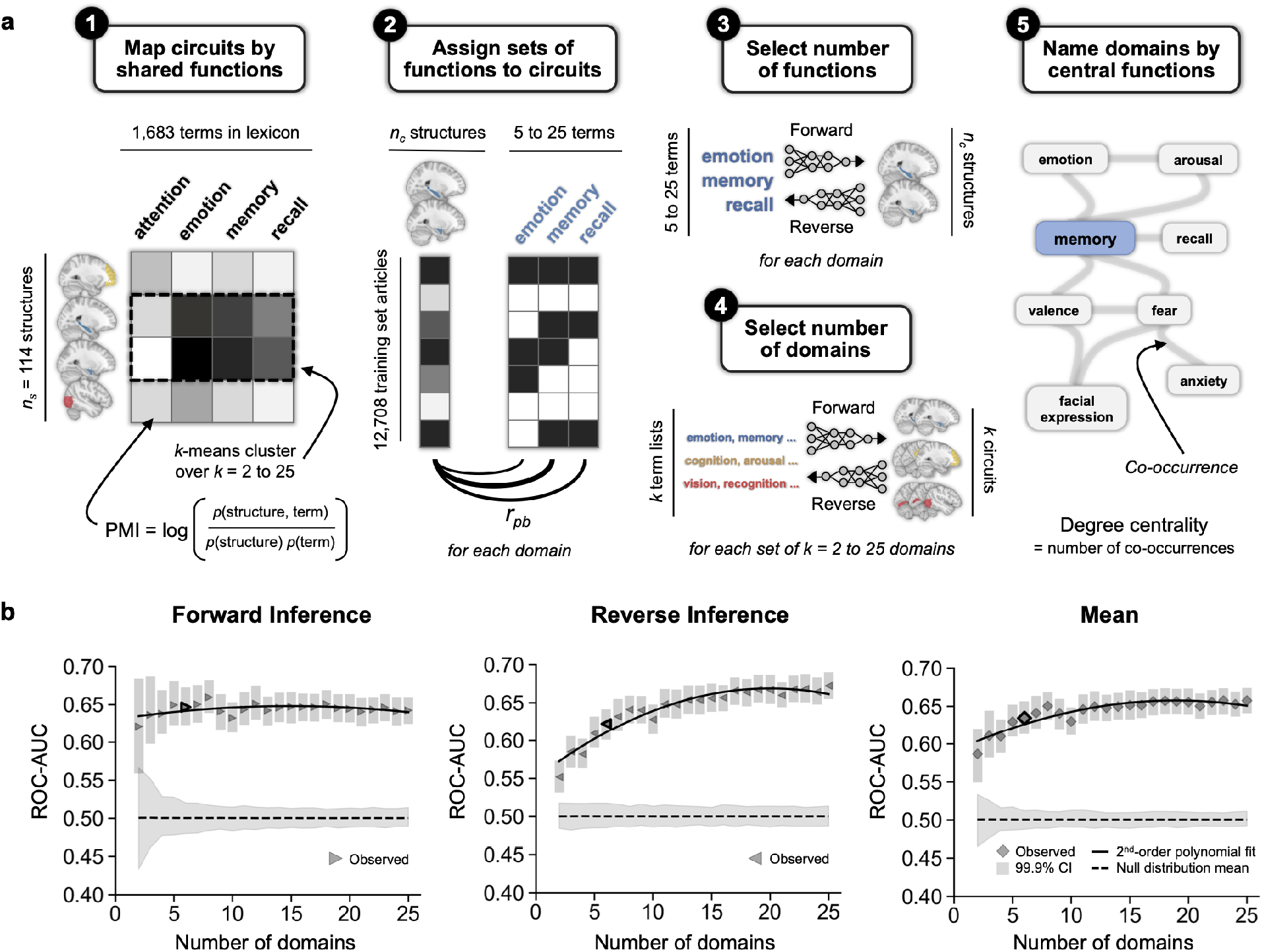
Approach to computational ontology. **a,** A data-driven ontology was generated in an integrative manner in the training set. First, 114 brain structures were clustered by *k*-means according to their cooccurrences with 1,683 terms for mental functions. The co-occurrence matrix was weighted by pointwise mutual information (PMI). Second, the top 25 terms for mental functions were assigned to each circuit based on the point-biserial correlation (*r_pb_*) of their binarized occurrences with the centroid of occurrences across structures. Third, the optimal number of terms was selected to maximize average ROC-AUC of logistic regression classifiers predicting structure occurrences from term occurrences (forward inference) and term occurrences from structure occurrences (reverse inference) over a range of term list lengths from 5 to 25. Fourth, the optimal number of domains was selected based on the average ROC-AUC of forward and reverse inference classifiers. Occurrences were summed across terms in each list and structures in each circuit, then thresholded by their mean across articles. In the fifth and final step, each domain was named by the mental function term with highest degree centrality of co-occurrences with other terms in the domain. **b,** Validation set ROC-AUC for classifiers used to select the optimal number of domains for the data-driven framework. Performance is plotted for forward inference classifiers (right-pointing triangles), reverse inference classifiers (left-pointing triangles), and their average (diamonds). Markers are outlined in black for the *k* = 6 solution, which was selected for the data-driven ontology as it was the lowest *k* value to achieve an average ROC-AUC along the asymptote. Shaded areas around markers represent 99.9% confidence intervals computed by resampling validation set articles with replacement over 1,000 iterations. The dashed line represents the mean of null distributions generated by shuffling true labels for validation set articles over 1,000 iterations, and the surrounding shaded area is the 95% confidence interval.

Candidate domains of the data-driven ontology are generated through an unsupervised learning approach that takes into account insights from information theory. To map links between the 1,683 terms for mental functions and 114 brain structures in our neuroanatomical atlas, we computed their co-occurrences across the training set. Co-occurrence values were reweighted by pointwise mutual information (PMI), which captures the extent to which the joint probability of a term and a brain structure occurring in the same article deviates from the expected probability if the two occurred independently of one another. Because PMI weighting is probability based, it results in high values for the most strongly associated functions and structures regardless of the raw number of times they were observed in the corpus. For instance, while “face identification task” is infrequent in article texts and few coordinates are mapped to the left amygdala, their cooccurrence has a high PMI value because they were both observed in the same small subset of articles. Circuits supporting distinctive sets of mental functions were then defined through *k*-means clustering of the brain structures by their PMI-weighted co-occurrences with mental function terms over a range of *k* values from 2 to 25.

Following this, we sought to identify the mental functions best representative of each circuit, which were selected in a manner that would reflect prevalence rates across the literature. We note that this was necessary because PMI gives high weight to connections that are specific and not necessarily common. While the pattern of structure-function connections is appropriate for grouping structures into circuits, a given connection may replicate only a few times in the literature. For the left amygdala, for instance, none of the top 25 terms with the strongest PMI weighting occurred in more than 0.2% of articles. We thus chose to instead assign the top 25 mental function terms to each circuit based on associations across the training set, computed as point-biserial correlations between binary term occurrences and the centroid of occurrences across structures in each circuit. For the circuit containing the left amygdala, the most strongly associated terms were “fear”, “emotion”, and “memory,” which occurred in 10.82%, 18.12%, and 17.74% of articles, respectively.

Next, the number of terms per data-driven domain is optimized through a supervised learning strategy. While up to 25 terms were initially assigned to a given circuit, fewer terms may suffice in representing its functional repertoire. In order to identify the set of terms and structures with the strongest predictive relationships, the number of mental function terms per circuit was determined by how well term occurrences predicted and were predicted by occurrences of structures. For each circuit, logistic regression classifiers were fit on the training set to predict structure occurrences from term occurrences (i.e., forward inference models), and to predict term occurrences from structure occurrences (i.e., reverse inference models), over a range of 5 to 25 mental function terms. The optimal number of mental function terms for each circuit was selected to maximize area under the receiver operating characteristic curve (ROC-AUC) averaged between the forward and reverse inference models in the validation set. The resulting term lists and the brain circuits mapped in the prior step can be examined interactively over *k* = 2 to 25 domains at http://nke-viewer.org/.

Finally, the optimal number of domains was established by evaluating forward and reverse inference classification performance in a similar fashion to the previous step. In this case, classifiers were designed to predict structure occurrences combined within circuits (forward inference) and term occurrences combined within the optimized word lists (reverse inference). Models were again evaluated in the validation set, with performance metrics averaged between forward and reverse inference models at each level of *k* (Fig. 1b). The resulting data-driven ontology comprises *k* = 6 domains with non-overlapping circuits that span the human brain and are associated with mental constructs consisting of 5 to 25 mental function terms. In the last step, each domain is named by the mental function term with highest degree centrality of its term-term co-occurrences. Examining the results for solutions with fewer domains reveals the early emergence of a limbic circuit supporting emotional/memory processes as well as a cortical circuit for cognitive functioning (Extended Data Fig. 4). These domains remain largely stable across increasing *k* values, with fractionation of subsets of brain structures into circuits for memory subprocesses, sensory perception, and language (Extended Data Fig. 5). We note that qualitatively similar results were obtained when using neural network classifiers instead of logistic regression to select the number of terms per domain and number of domains in total (Extended Data Fig. 6-8).

Expert-determined frameworks for basic brain function (i.e., RDoC) and psychiatric illness (i.e., the DSM) were mapped in a top-down fashion beginning with their terms for mental function and dysfunction. NLP was applied to translate the language of the frameworks into the language of the human neuroimaging literature, and the resulting term lists were subsequently mapped onto circuits of co-occurring brain structures (Fig. 2a). For RDoC, word embeddings were trained using GloVe on a corpus of 29,828 articles (Extended Data Fig. 1b) that was curated by supplementing the original corpus with the results of a PubMed query for human neuroimaging studies (Extended Data Fig. 3a). For the DSM, embeddings were trained with the same parameters on a corpus of 26,070 articles (Extended Data Fig. 1c) enriched with human neuroimaging studies of mental illness retrieved from PubMed (Extended Data Fig. 3b). Candidate synonyms were identified among terms for mental function or dysfunction (Extended Data Fig. 2) by the cosine similarity of their embeddings to the centroid of seed embeddings in each domain. This approach yielded synonyms of the RDoC domains with higher semantic similarity to seed terms than a previously published NLP approach^19^ (Fig. 2b). DSM domains were additionally required to contain terms that appeared in at least 5% of articles in the corpus with coordinate data. Finally, we mapped brain circuits from each term list based on PMI-weighted co-occurrences with brain structures across the full corpus of articles with coordinates (*n* = 18,155 articles), restricting the circuits to positive values with FDR < 0.01. Altogether, RDoC and DSM domains each comprise 5 to 25 terms for mental function/dysfunction and a circuit of co-occurring brain structures.

**Fig. 2.**
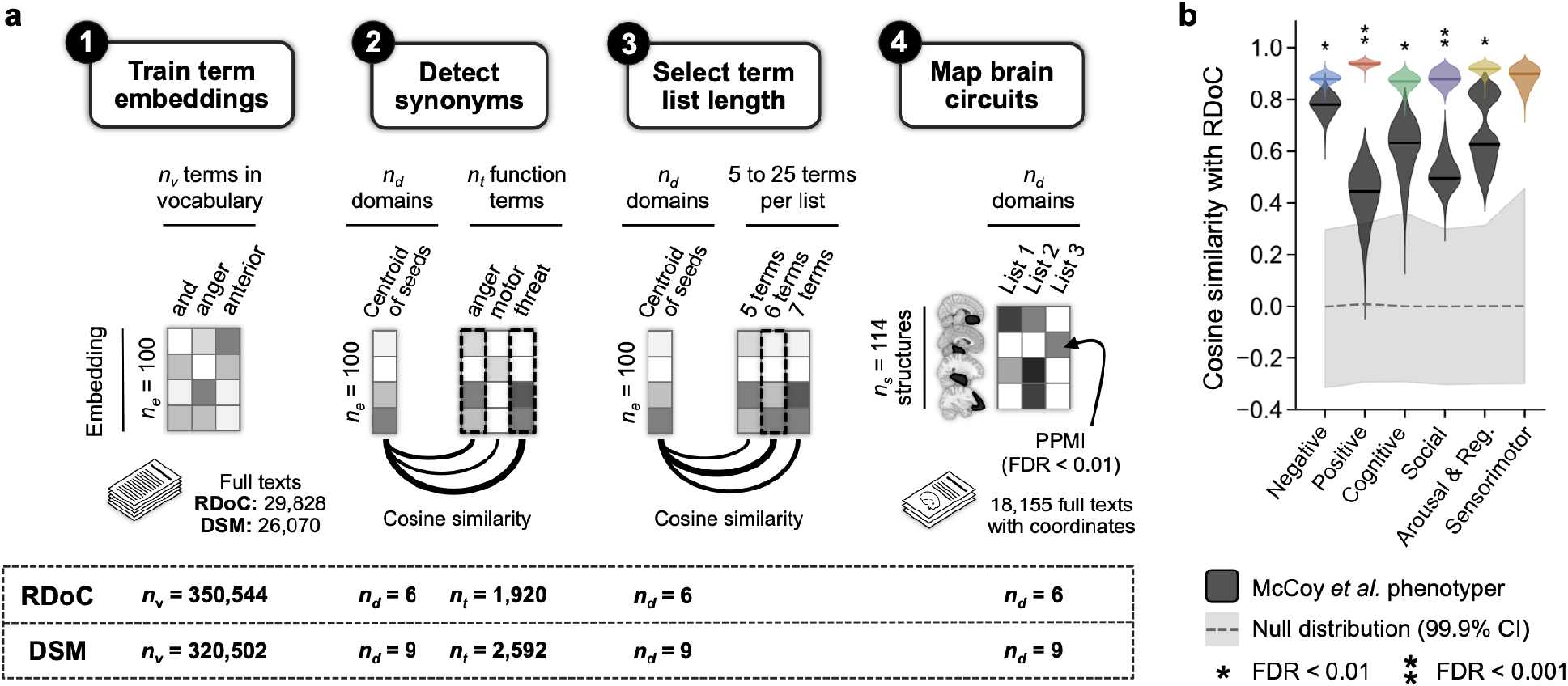
Approach to mapping expert-determined frameworks for brain function (RDoC) and mental illness (DSM). **a,** Seed terms from the RDoC and DSM frameworks were translated into the language of the human neuroimaging literature through a computational linguistics approach. First, word embeddings of length 100 were trained using GloVe. For RDoC, embeddings were trained on a general human neuroimaging corpus of 29,828 articles (Extended Data Fig. 1b). For the DSM, embeddings were trained on a psychiatric human neuroimaging corpus of 26,070 articles (Extended Data Fig. 1c). Candidate synonyms included terms for mental functions in the case of RDoC and for both mental functions and psychopathology in the case of the DSM, as detailed in Extended Data Fig. 2. In the second step, the closest synonyms of seed terms were identified based on the cosine similarity of synonym term embeddings with the centroid of embeddings across seed terms in each domain. Third, the number of terms for each domain was selected to maximize cosine similarity with the centroid of seed terms. The mental function term lists for each domain were then mapped onto brain circuits based on positive pointwise mutual information (PPMI) of term and structure co-occurrences across the corpus of 18,155 articles with activation coordinate data (Extended Data Fig. 1a). Structures were included in the circuit if the FDR of the observed PPMI was less than 0.01, determined by comparison to a null distribution generated by shuffling term list features over 10,000 iterations. **b,** Semantic similarity to seed terms in the RDoC framework for our term lists generated using GloVE (colored) compared to a baseline from the literature (dark gray). The baseline model includes term lists generated by McCoy *et al.* through latent semantic analysis followed by a filtering procedure.^19^ Bootstrap distributions for each domain were generated by resampling the 100-*n* embedding dimension with replacement over 10,000 iterations, then assessed for a difference in means (* FDR < 0.01, ** FDR < 0.001). Solid lines denote the observed similarity values. No comparison is shown for Sensorimotor Systems, which was added to RDoC following publication of the McCoy *et al.* phenotyper. Null distributions were generated for the GloVe term lists by shuffling embeddings over 10,000 iterations. Gray dashed lines denote the null distribution means.

A comparison of the data-driven ontology with the expert-determined frameworks reveals notable differences in circuit-function mappings (Fig. 3). First, the data-driven ontology offers novel combinations of emotional and cognitive terms in its domains for *Memory* and *Reaction Time*, which each relate to several domains in the RDoC and DSM frameworks. Likewise, the RDoC domain for *Cognitive Systems* relates strongly to both *Manipulation and Vision* in the data-driven ontology, indicating that further functional specification may be warranted in RDoC. While the *Reward* domain of the data-driven framework is similar to a single RDoC domain for *Positive Valence* at the FDR < 0.01 threshold, the *Reward* circuitry is defined more specifically by frontomedial regions and the nucleus accumbens. Relations between the data-driven ontology and the DSM are sparser overall, with four of the six data-driven domains exhibiting above-threshold similarity with four of the nine DSM domains. *Memory* in the data-driven ontology is similar to *Anxiety* and *Trauma and Stressor* domains (but not *Bipolar* and *Depressive* domains with asymmetric fronto-subcortical circuitry). The *Developmental* domain in the DSM maps onto three cortical domains of the data-driven ontology, including the *Manipulation* domain that also has high similarity with *Psychotic* illness.

**Fig. 3.**
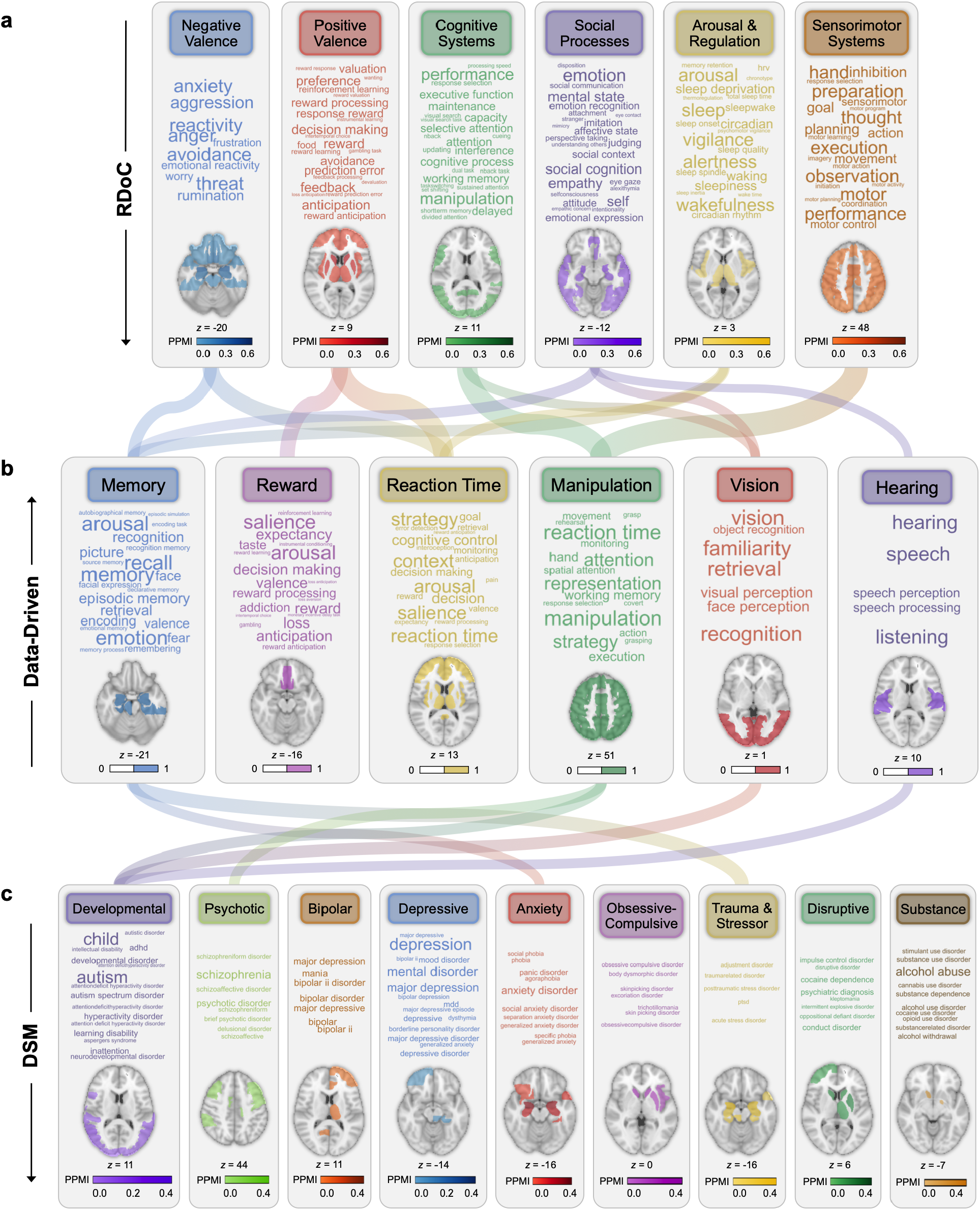
Data-driven ontology of brain functions related to expert-determined frameworks for brain function (RDoC) and mental illness (DSM). Links are scaled to the Dice similarity of mental function terms and brain structures in each domain (FDR < 0.01 based on permutation testing over 10,000 iterations). Word size is scaled to frequency in the corpus of 18,155 articles with activation coordinate data. **a,** The RDoC framework was modeled in a top-down manner from terms to brain circuits (Fig. 2a). **b,** A data-driven ontology for domains of brain function was engineered in a bottom-up manner beginning with circuits (Fig. 1a). **c,** The DSM framework for mental disorders was modeled in a top-down manner by a procedure analogous to that for RDoC (Fig. 2a).

We next sought to evaluate our data-driven ontology and the expert-determined frameworks with respect to three organizing principles of key relevance to neuroscience: reproducibility, modularity, and generalizability. Reproducibility concerns whether the circuit-function links underlying domains are well predicted from their observed co-occurrences in the neuroimaging literature. Human neuroimaging has demonstrated that several brain regions (e.g., the insula and anterior cingulate) are widely activated across task contexts, rendering them unreliable predictors of mental state.^15, 20^ If links between brain circuits and mental functions are not reproducible across studies, then the ontological entities and neuropsychiatric biomarkers derived from them will be of limited utility. We assessed the reproducibility of circuit-function links based on the performance of logistic regression classifiers predicting the occurrence of mental function terms in article text from coordinate data mapped to the structures of our neuroanatomical atlas (Figure 4a). Binary scores for the mental functions listed under each domain were computed by mean-thresholding term occurrences, then mean-thresholding the sum of terms within each domain list. Hyperparameters were tuned on the validation set of 20% of articles, and classifiers were evaluated by ROC-AUC in the test set containing 10% of articles (*n*=1,816; Figs. 4b-g). ROC-AUC was higher across domains of the data-driven ontology as compared to RDoC and the DSM (Fig. 4h). Whereas all domains of the data-driven and RDoC frameworks achieved above-chance ROC-AUC, the *Psychotic* domain in the DSM framework did not (Fig. 4f). These results suggest that orienting neurobiological and psychiatric frameworks around the circuits and term lists derived through our data-driven approach could improve the reproducibility of their structure-function links. Further results supporting this conclusion were obtained when evaluating classifiers by F1 score (Extended Data Fig. 9), when using mental function terms to predict occurrences of brain structures (Extended Data Figs. 10-11), and when repeating analyses with neural network classifiers (Extended Data Figs. 12-18).

**Fig. 4.**
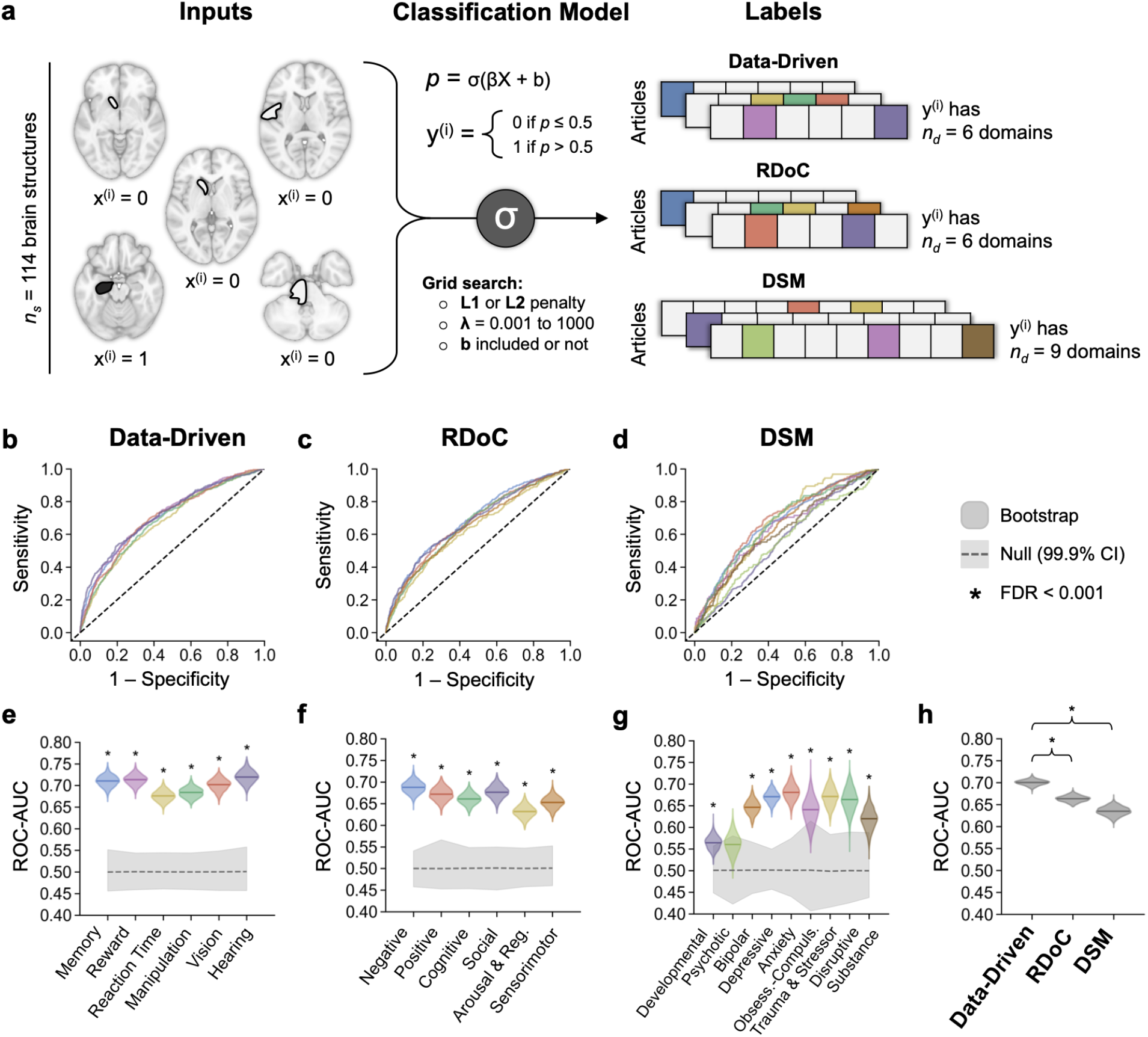
Mental functions defined in a data-driven manner have more reproducible links with locations of brain activity. **a,** Logistic regression classifiers were trained to predict the occurrence of terms for mental functions in neuroimaging article full texts from the brain activation coordinates that articles reported. Inputs included activation coordinate data mapped to 114 brain structures in a wholebrain neuroanatomical atlas. Labels were term occurrences thresholded by mean frequency across the corpus, then the mean frequency of terms in each domain. Classifiers were trained over 1,000 iterations with hyperparameters selected through a grid search over the following values: penalty = L1 or L2; regularization strength = 0.001, 0.01, 0.1, 1, 10, 100, or 1,000; intercept included or not. Training was performed in 70% of articles (*n* = 12,708), hyperparameters were tuned in 20% of articles, and classifiers were evaluated in a test set containing 10% of articles (*n* = 1,816). ROC curves are shown for the test set performance of classifiers with mental function features defined by **b,** the data-driven ontology, **c,** RDoC, and **d,** the DSM. **e-g,** Bootstrap distributions of ROC-AUC (colored) were computed by resampling articles in the test set with replacement over 1,000 iterations. Observed values in the test set are shown with solid lines. Null distributions (gray) were computed by shuffling true labels for term list scores over 1,000 iterations; the 99.9% confidence interval is shaded, and distribution means are shown with dashed lines. The mean of each bootstrap distribution was assessed for a difference in mean from its corresponding null distribution (* FDR < 0.001). **h,** ROC-AUC across the domains in each framework. Solid lines denote means of the bootstrap distributions macro-averaged across classifiers. Differences in bootstrap means were assessed for each framework pair (* FDR < 0.001).

The second organizing principle of interest in constructing an ontology of brain function is modularity – namely, the extent to which domains are internally homogeneous and distinctive from one another in their patterns of functions and structures. The principle of modulatory has been observed across neural measures and scales, ranging from single neurons in *C. elegans*^21^ to distributed resting-state fMRI networks in humans. ^22^ However, because task-based neuroimaging studies are limited in the number of mental states they can reasonably induce, it is largely unknown whether task-related brain activity is similarly modular. Our automated meta-analytic approach enables us to overcome this limitation to the extent one can assume that articles reporting different mental constructs and brain structures in their texts and data are studying different underlying domains of brain function. We began by assigning articles to the most similar domain of each framework, yielding “subfields” of human neuroimaging (Fig. 5a-c; Extended Data Fig. 19). Consistent with high comorbidity rates^23^ and similar neural alterations^24^ between affective disorders, there is visible overlap among the *Bipolar*, *Depressive*, and *Anxiety* illness domains of the DSM (Fig. 5c). Modularity was then assessed by the ratio of mean Dice distance of articles between versus within subfields (Fig. 5d-f). The domain-level results exceeded chance for all domains across the three frameworks. Macro-averaging across domains in each framework, we find that modularity is higher for the data-driven ontology compared to both RDoC and the DSM (Fig. 5g). Surprisingly, modularity of the DSM domains exceeded that of the RDoC domains. These results support the movement currently underway to ground psychiatric diagnoses in brain circuits for transdiagnostic mental constructs,^13, 25^ while at the same time cautioning against the assumption that expert-determined domains of brain function will lead to improved ontological modularity.

**Fig. 5.**
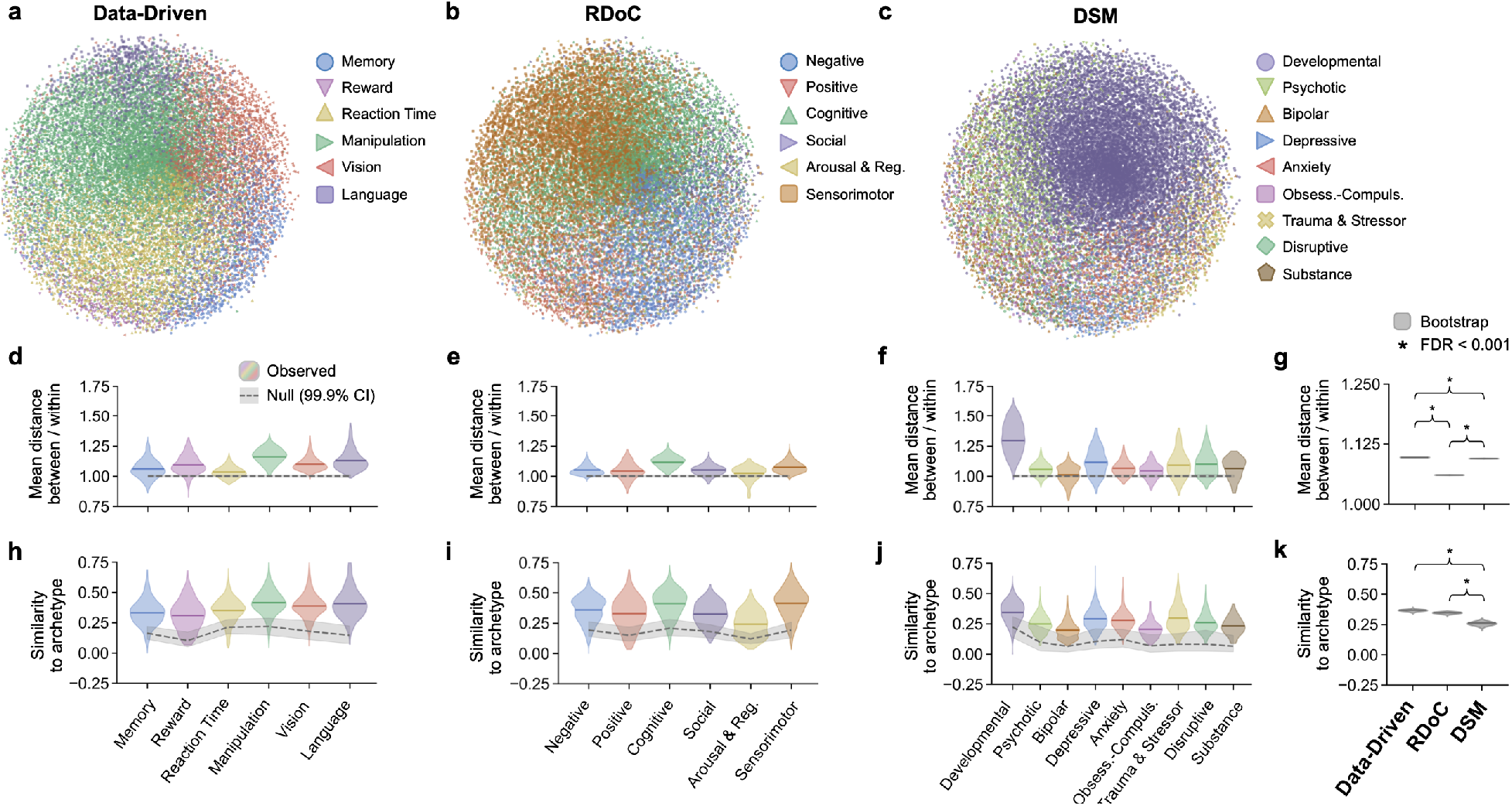
The data-driven ontology partitions the neuroimaging literature into modular subfields, for which domains are generalizable representations of brain circuits and mental functions. Articles were matched to domains based on the Dice similarity of mental function terms and brain structures. **a-c,** Multidimensional scaling of Dice distances between articles colored by domain assignments. Articles are colored by their domain assignments. **d-f,** Modularity of the article partitioning was assessed by comparing the mean Dice distance of function and structure occurrences of articles between domains versus within domains. Observed values are colored by domain; null distributions in gray were computed by shuffling distance values across article partitions. **g,** Bootstrap distributions of modularity macro-averaged across the framework domains were assessed for differences in means between frameworks (* FDR < 0.001). **h-j,** Generalizability was assessed by computing the Dice similarity of each domain’s “archetype” vector of function terms and brain structures with the terms and structure occurring in each article of the domain’s partition. Observed values are colored by domain; null distributions in gray were computed by shuffling terms and structures in each archetype. **k,** Bootstrap distributions of generalizability macro-averaged across domains were assessed for differences in means between frameworks (* FDR < 0.001). All bootstrap and null distributions shown were generated over 1,000 iterations.

The third principle of central relevance to an ontology of brain function is generalizability. By this principle, the pattern of functions and structures included in each domain of the ontology should be a representative archetype of the functions and structures occurring in single articles, and presumably, in the underlying neurobiological phenomena they address. Previous metaanalyses have demonstrated that some (though not all) psychological domains have generalizable representations in the activity of specialized brain regions.^26, 27^ We tested generalizability by computing the similarity of function and structure occurrences in each article to the archetypal function-structure pattern of the domain to which it was assigned (Figs. 5h-j). Similarity to the archetype exceeded chance for all frameworks tested, supporting the interpretation that they represent information which generalizes well within the subfields of human neuroscience. Yet, further gains in similarity to the archetype across domains were achieved by both the data-driven ontology and RDoC relative to the DSM (Fig. 5k), highlighting the disconnect between current understanding of brain function and the way that mental disorders have historically been categorized. We anticipate that if mental disorders were redefined as disruptions in basic brain circuitry, their information content would better generalize within subfields of the human neuroscience literature.

As human neuroscience expands beyond what any individual can reasonably interpret, the field demands computational approaches to ontology that have been developed in biomedical informatics.^2^ We demonstrate here that it is feasible to synthesize the results of thousands of articles through a combination of text mining and machine learning techniques, organizing the knowledge of the human neuroimaging discipline into domains jointly defined by mental constructs and brain circuitry. Our procedure for modeling domains of brain function can be readily adapted to summarize other neuroscience subfields, such as direct electrophysiological recordings, cellular-level imaging, or tract tracing experiments. It will be important to gather further evidence of convergent validity for the domains of the data-driven ontology as the results of our meta-analysis may in part reflect the spatial and temporal limitations of MRI and PET. For instance, while the data-driven ontology defines a single brain circuit for processing emotional memory, neuronal population recordings in the rodent amygdala reveal functional divisions for positive and negative affective valence.^28^ Nonetheless, our meta-analytic approach has the advantage of accounting for functional similarities and differences on the whole-brain scale in human subjects.

The comparative analyses we undertook here underscore the need for rigorous ontological standards in the neuroscience and psychiatry research communities, as well as for flexible methods to hold such standards against ontologies built through varying approaches. Our assessments of reproducibility, modularity, and generalizability indicate that the data-driven ontology meets criteria for predictive, discriminant, and convergent validity, respectively. We conclude that the data-driven ontology, with its novel mappings of affective and cognitive processes, represents a robust framework for conceptualizing brain function. RDoC likewise upheld the ontological standards assessed here, though gains in reproducibility and modularity metrics were achieved with the data-driven organization of knowledge. The DSM, by contrast, failed to uniformly achieve predictive validity across its domains. Future work may seek to refine the data-driven ontology by applying techniques aimed at explicitly measuring and controlling for quantifiable biases. Neuroscience ontologies can be further leveraged if they are made publicly available in a structured form that is amenable to automated hypothesis generation.^29^ Going forward, we expect that automated approaches to ontology will pave the way for a new science of synthesis able to accelerate progress in a wide range of disciplines.

## Methods

### Neuroimaging Corpora

Three corpora of neuroimaging articles were curated from existing databases and by web scraping (Extended Data Fig. 1). All articles were manually inspected by EB to ensure they conducted neuroimaging of the human brain using MRI, PET, or other standard techniques.

#### Neuroimaging Corpus with Coordinate Data

First, a corpus of brain activation coordinates and neuroimaging article full texts (*n* = 18,155; Extended Data Fig. 1a) was collected as the substrate for the data-driven ontology (Fig. 1a) and for coordinate-based analyses (Figs. 4, 5). Articles were included if they reported coordinates in the human brain in standard Montreal Neurological Institute (MNI) or Talairach space. Coordinates were gathered on the study level from BrainMap^**Error! Bookmark not defined.**^ (*n* = 3,346), then Neurosynth^15^ (*n* = 12,676), and finally by deploying the Automated Coordinate Extractor (*n* = 2,133).^30^

#### General Neuroimaging Corpus

A comprehensive corpus of neuroimaging article full texts (*n* = 19,828; Extended Data Fig. 1b) served as the basis for our computational linguistics approach to selecting mental function terms for RDoC (Fig. 2a). Articles were retrieved in response to a PubMed query (Extended Data Fig. 3a) and combined with the first corpus.

#### Psychiatric Neuroimaging Corpus

A corpus of human neuroimaging articles enriched with studies of psychiatric illness (*n* = 26,070; Extended Data Fig. 1c) served as the basis for selecting mental function and dysfunction terms for the DSM. Articles were retrieved from PubMed through a query for DSM-5 disorders (Extended Data Fig. 3b) and combined with those from the first corpus. A characterization of key terms appearing in article full texts over time reveals that, as expected, the term *DSM* occurs in a consistently high proportion of articles in the psychiatric corpus (Extended Data Fig. 1f) relative to general neuroimaging corpora (Extended Data Figs. 1d, 1e).

### Coordinate Processing

Brain activation coordinates were converted to MNI space if reported in Talairach using the Lancaster transform.^31^ FSL atlasquery (version 5.0.10) mapped coordinates to 114 bilateral gray matter structures in probabilistic anatomical atlases of the human cerebrum^17^ and cerebellum.^18^ Structures were retained if their probability of containing the coordinate exceeded zero, resulting in one-to-many mappings from each coordinate to the brain. Differences in reporting practices were mitigated by binarizing the mappings for each study, meaning that a study could not report data in a given structure more than once.

### Natural Language Processing

#### Text Preprocessing

Article full texts were extracted from PDF or HTML files downloaded through Stanford University subscription services, from PDF files available through the PMC Open Access Subset, or from XML files in the PMC Author Manuscript Collection. In order to tokenize articles, a lexicon was compiled from public sources (Extended Data Fig. 2). For RDoC and the data-driven ontology, the lexicon was limited to terms describing mental processes and the paradigms used to study them. The lexicon for the DSM was extended to terms for psychopathology. Preprocessing of texts and the lexicon included case-folding, removal of stop words and punctuation, and lemmatization with WordNet. *N*-grams listed in the lexicon were combined with underscores in article texts.

#### Word Embeddings

To identify synonyms of seed terms in existing frameworks as shown in Fig. 2a, word embeddings were trained with GloVe^32^ on concatenated full texts in the general neuroimaging corpus (for RDoC) and psychiatric neuroimaging corpus (for the DSM). The GloVe parameters were as follows: embedding dimension = 100, minimum word count = 5, window size = 15 words, iterations = 500.

### Computational Ontology

#### Data-Driven Ontology

The data-driven ontology was built up from relationships between (1) activation coordinate data mapped to brain structures and (2) terms for mental functions in article full texts. To prevent over-estimation of classifier test set performance, the data-driven ontology was generated in a training set that contained 70% of articles (*n* = 12,708). Co-occurrences were computed between brain structures (*n_s_* = 114) and terms for mental functions (*n_t_* = 1,683) across training set articles, with re-weighting by PMI to reduce the effects of structure and term frequency. PMI was computed as the logarithm of the observed co-occurrence value divided by the expected co-occurrence value. To avoid taking the logarithm of zero, terms and structures with no cooccurrences were assigned a value of zero. Circuits were then mapped by *k*-means clustering of brain structures by PMI-weighted links with mental function terms over a range of *k* values from 2 to 25. Up to 25 terms for mental functions were assigned to each circuit by the point-biserial correlation (*r_pb_*) of the centroid of occurrences of the circuit’s structures with binary function term occurrences. The optimal number of circuits and terms for each circuit was determined based on the validation set performance of classifiers described below under *Classification Models: Data-Driven Optimization*. The resulting circuits and lists of their associated terms are what we term “domains.” Finally, each domain was named by the mental function in its list with highest degree centrality of term-term co-occurrences across training set articles.

#### RDoC and DSM Frameworks

RDoC and the DSM were modeled by translating their terms for mental functions and disorders into the language of the human neuroimaging literature, then mapping the resulting term lists to circuits of brain structures co-occurring across the literature (Fig. 2a). RDoC seed terms were preprocessed from RDoC behaviors, self-report items, and paradigms grouped by domain; DSM seed terms were preprocessed from disorders grouped by the headings of Section II in Edition 5 (which we refer to subsequently as “domains”). Candidate synonyms from the lexicon (Extended Data Fig. 2) were identified by cosine similarity to the centroid of seed terms in each domain over a range of list lengths from 5 to 25 terms. The list that maximized cosine similarity to the centroid of seed terms was selected for each domain. For the DSM, to reduce overlap between frequently co-occurring disorders, seed terms were additionally required to be unique to domains and candidate synonyms to have a cosine similarity ≥ 0.5. Co-occurrences weighted by positive PMI (PPMI) were then mapped between brain structures and term lists across the corpus of articles with coordinate data (*n* = 18,155). Brain circuits were defined for each domain by retaining structures with PPMI values and FDR < 0.01 based on comparison to a null distribution generated over 10,000 iterations of shuffling term list occurrences across articles.

### Logistic Regression Classifiers

Classifiers were designed to predict the locations of brain activation coordinates reported by articles from the terms for mental functions occurring in article full texts (i.e., forward inference), and in turn, to predict term occurrences from coordinate locations (i.e., reverse inference). Articles with coordinate data were split into sets for training (70%; *n* = 12,708), validation (20%; *n* = 3,631) and test (20%; *n* = 1,816). All classifiers were fit in the training set with the following hyperparameters optimized over a grid search to maximize validation set ROC-AUC: regularization penalty (L1 or L2), regularization strength (0.001, 0.01, 0.1, 1, 10, 100, or 1,000), and intercept (included or not). The classifiers used in optimizing the number of terms per data-driven domain (Fig. 1a, Step 3) were fit over 100 iterations, and those used in optimizing the number of circuits were fit over 500 iterations (Fig. 1a, Step 4). The test set was reserved for the final comparison of the data-driven ontology with RDoC and the DSM, each of which scored article texts using a different list of terms for mental functions (see *Reproducibility*). The classifiers used in reproducibility analyses were optimized over the hyperparameter grid described above with training for 1,000 iterations.

### Neural Network Classifiers

All neural network classifiers were fit in PyTorch with the Adam optimizer (β_1_ = 0.9, β_1_ = 0.999, ε = 10^-8^). The architecture always included 8 fully connected (FC) layers. Batch normalization and a rectified linear unit (ReLU) activation function were applied to the first seven layers, with dropout during training. A sigmoid activation function was applied to the last layer for one-versus-all classification. The classifiers used in optimizing the number of terms per data-driven domain were fit with learning rate = 0.001, weight decay = 0.001, neurons per layer = 100, dropout probability = 0.1, batch size = 1,024, epochs = 100. The classifiers used to select the number of data-driven domains (Extended Data Fig. 6a) were fit with the same hyperparameters over 500 epochs. Classifiers used in the comparison between the data-driven, RDoC, and DSM frameworks (Extended Data Figs. 12-15) were optimized through a randomized grid search over 100 combinations of the following hyperparameters: learning rate (10^-5^ to 1 on a log scale), weight decay (10^-5^ to 1 on a log scale), number of neurons per layer (25 to 150 by 25), and dropout probability (0.1 to 0.9 by 0.1). Each classifier was trained for 500 epochs in this grid search, and the optimal hyperparameter combination was selected to maximize ROC-AUC evaluated on the validation set. Classifiers were subsequently trained for 1,000 epochs with the selected hyperparameters. The test set was reserved for the final comparison of the data-driven ontology with RDoC and the DSM, each of which scored article texts using a different list of terms for mental functions.

#### Data-Driven Optimization

The data-driven ontology was built through the computational meta-analyses outlined in Fig. 1a, which used classifier performance to select thresholds at two stages. First, after mapping brain circuits and associating them with terms for mental functions, classifier performance determined the optimal number of terms per circuit. Specifically, classifiers were trained over a range of term list lengths (5 to 25 terms) and number of circuits (2 to 25). Forward inference classifiers predicted occurrences of coordinates mapped to structures in each circuit from occurrences of terms in its corresponding list; reverse inference classifiers predicted term occurrences from structure occurrences. For each circuit, the number of terms was chosen to maximize the mean ROC-AUC of forward and reverse inference classifiers applied to the validation set.

Second, after assigning terms to each circuit, classifier performance determined the optimal number of circuits. Classifiers were trained again over the range of circuit numbers (2 to 25), this time using occurrences grouped by term list and circuit. Forward inference classifiers predicted occurrences of structures summed by circuit from occurrences of terms summed by list across the full set of domains; reverse inference classifiers predicted term list occurrences from circuit occurrences, again across the full set of domains. Input and output data were binarized by mean values across articles. The optimal number of domains was chosen to maximize the mean ROC-AUC of forward and reverse inference classifiers applied to the validation set.

#### Reproducibility

Test set performance was compared for classifiers that based their inputs or outputs on term lists for each domain of the data-driven ontology, RDoC, and the DSM. Term lists were used to score article full texts through the following process: (1) term occurrences were thresholded by their mean across articles, (2) the binarized term occurrences were summed across terms in each list, (3) the sums were thresholded by their mean across articles. Neural data for classification models included the binarized occurrences of coordinates within 114 brain structures. Reverse inference classifiers took brain structure occurrences as inputs to predict text scores as outputs (Fig. 4); conversely, forward inference classifiers took the text scores as inputs to predict brain structure occurrences as outputs (Extended Data Fig. 10). Performance was assessed for each classifier based on overlap of bootstrap and null distributions for ROC-AUC computed in the test set of articles. Bootstrap distributions were computed by resampling articles in the test set with replacement over 1,000 iterations. Null distributions were computed by shuffling true labels for term lists (Fig. 4) or brain structures (Extended Data Fig. 10) over 1,000 iterations.

### Article Partitioning

The data-driven ontology, RDoC, and the DSM were applied to partition articles across the corpus (*n* = 18,155) into subfields of neuroscience. Each domain was represented by a binary vector with one-valued cells for the term and brain structures it entails, as shown in Fig. 3. Likewise, each article was represented by a binary vector with one-valued cells for the terms that occurred in its full text and the brain structures that were mapped from its reported activation coordinates. Articles were assigned to the domain in each framework with the highest Dice similarity of its terms and structures, as shown in Figs. 5a-c and Extended Data Fig. 19.

#### Multidimensional Scaling

The Dice distance between articles across the corpus (*n* = 18,155) was visualized with multidimensional scaling (Figs. 5a-c). Distances were computed between articles based on the binary vectors for the terms and structures they reported. Metric multidimensional scaling was performed on the square distance matrix (18,155 x 18,155) with epsilon = 0.001 over 5,000 iterations. The resulting matrix (18,155 x 2) was visualized with articles colored by their domain assignments in the data-driven ontology (Fig. 5a), RDoC (Fig. 5b), and the DSM (Fig. 5c).

#### Modularity

Modularity of the partitions was assessed by the ratio of mean Dice distance of articles between versus within domain partitions (Figs. 5d-f). Dice distance was computed over the binary mental function term and brain structure occurrences in each article. Null distributions were computed by shuffling the distance matrix over 1,000 iterations. Bootstrap distributions were computed by resampling distances with replacement over 1,000 iterations (Fig. 5g).

#### Generalizability

Generalizability of each domain to articles within its corresponding partition was assessed by Dice similarity of terms and structures (Figs. 5h-j). Null distributions were computed by shuffling terms and structures in each domain over 1,000 iterations. Bootstrap distributions were computed by resampling articles with replacement over 1,000 iterations (Fig. 5k).

### Statistics

Unless otherwise noted, statistical significance of observed values was determined by testing the null hypothesis they did not differ from the mean of corresponding distributions of shuffled data. Frameworks were compared by testing for a difference in means of bootstrap distributions. FDR estimates were computed by correcting *p*-values according to the Benjamini-Hochberg method.^33^

## Code Availability

The code used to generate and assess knowledge frameworks is available at https://github.com/ehbeam/neuro-knowledge-engine. The code supporting the interactive viewer for the data-driven ontology can be accessed at https://github.com/ehbeam/nke-viewer.

