## Extended Data for "A computational knowledge engine for human neuroscience"

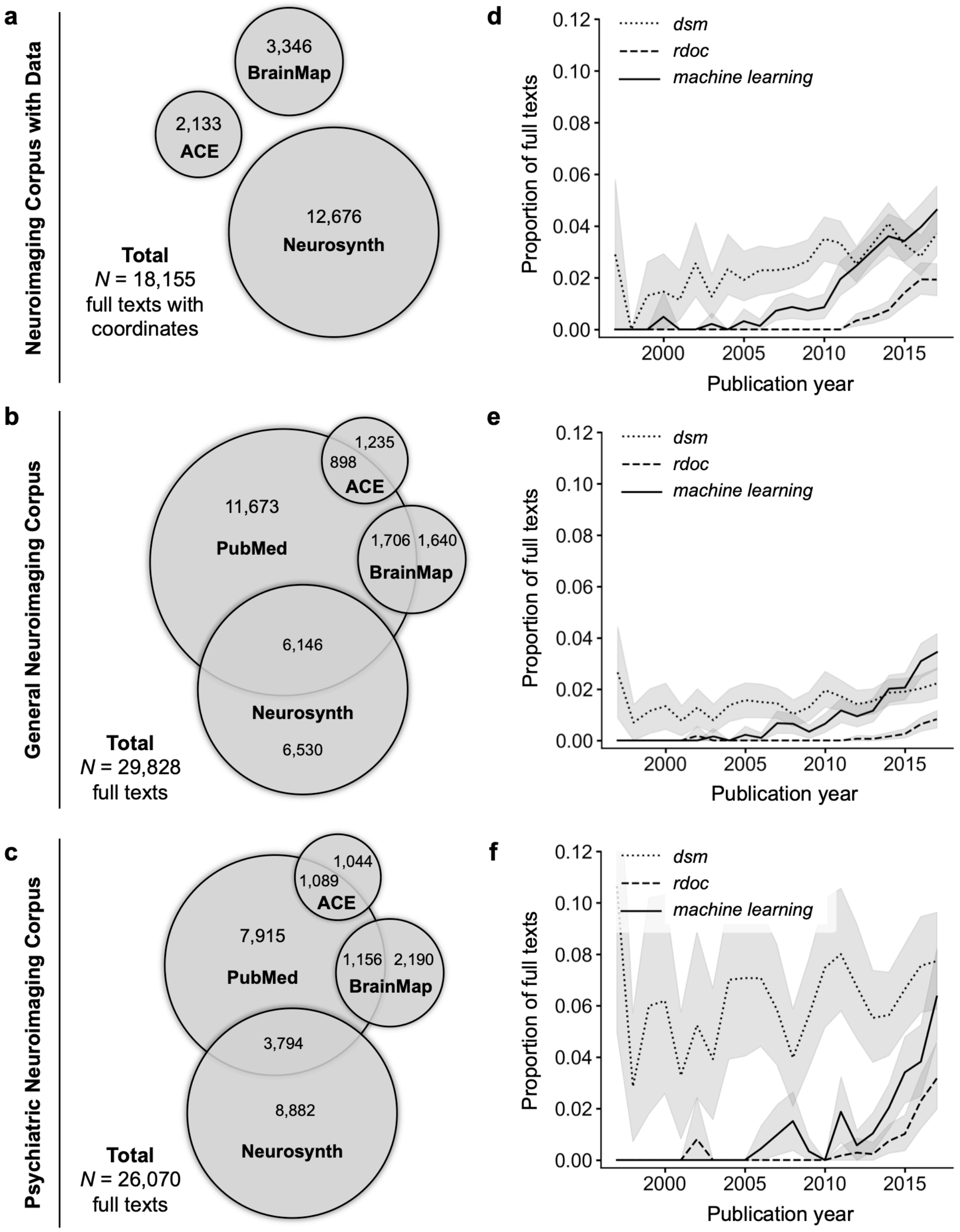


**Extended Data Fig. 1 | Corpora of human neuroimaging articles. a,** Articles reporting locations of activity in the human brain in standard MNI or Talairach space. Article metadata and coordinates were curated first from BrainMap (*n* = 3,346), then from Neurosynth (*n* = 12,676), then by deploying the Automated Coordinate Extractor (*n* = 2,133).[1] **b,** A comprehensive corpus of human neuroimaging articles served as the basis for a computational linguistics approach to selecting mental function words for the RDoC framework. Articles were retrieved in response to a PubMed query (Extended Data Fig. 3a) and combined with those reporting coordinate data. **c,** A corpus of human neuroimaging articles enriched with studies addressing psychiatric illness served as the basis for selecting mental function and dysfunction words for the DSM framework. As before, articles were retrieved through a PubMed query (Extended Data Fig. 3b) and combined with those reporting coordinate data. **d-f,** Characterizations of the corpora by the proportion of article full texts in which the terms *dsm*, *rdoc*, and *machine learning* occurred at least once, broken down by year. As expected, the proportion for *dsm* is consistently high over time in the psychiatric corpus. The proportion for *rdoc* begins increasing in 2010 when the first working group was held to populate the framework. Its proportion is lower than that of *machine learning*, which has increased since the mid-2000s. Shaded areas show bootstrap distributions generated by resampling articles published in a given year with replacement over 1,000 iterations.

| Ontology | Entities | Data-Driven | RDoC | DSM | Source |
|---|---|---|---|---|---|
| BrainMap | ○ Behavioral Domains<br>○ Paradigm Classes | ✔ | ✔ | ✔ | brainmap.org/taxonomy |
| Cognitive Atlas | ○ Behaviors<br>○ Concepts<br>○ Personality Traits<br>○ Tasks | ✔ | ✔ | ✔ | cognitiveatlas.org |
| | ○ Disorders | | | ✔ | |
| Cognitive Paradigm Ontology | ○ Stimulus Modality<br>○ Explicit Stimulus<br>○ Stimulus Role<br>○ Response Modality<br>○ Overt Response<br>○ Instructions<br>○ Paradigm Classes | ✔ | ✔ | ✔ | wiki.cogpo.org |
| Diagnostic & Statistical Manual of Mental Disorders | ○ Axis I & II Diagnostic Classifications<br>○ ICD-9-CM & ICD-10-CM | | | ✔ | DSM-III, DSM-IV-TR, DSM-5 |
| Medical Subject Headings | ○ Psychiatry & Psychology (F) | | | ✔ | ncbi.nlm.nih.gov/mesh |
| Mental Functioning Ontology | ○ Mental Functions | ✔ | ✔ | ✔ | github.com/jannahastings/ mental-functioning-ontology |
| | ○ Mental Disorders | | | ✔ | |
| Neuroscience Informatics Framework | ○ Functions | ✔ | ✔ | ✔ | github.com/SciCrunch/NIF-Ontology |
| Research Domain Criteria | ○ Behavior<br>○ Self-Report<br>○ Paradigms | | ✔ | | nimh.nih.gov/research/ research-funded-by-nimh/rdoc/ constructs/rdoc-matrix.shtml |

**Extended Data Fig. 2 | Mental function lexicon broken down by framework.** For the data-driven ontology, brain circuits were clustered by their PMI-weighted co-occurrences with the indicated terms, which were subsequently assigned to domains by the procedure detailed in Fig. 1a. For RDoC and DSM frameworks, terms in their respective lexicons were assigned based on semantic similarity to the centroid of seeds in each domain, as depicted in Fig. 2a.

**a** **General Neuroimaging Corpus**

```
((positron emission tomography) OR “fmri”) AND (psychiat* OR psychol* OR mental OR emotion* OR
reward* OR cognit* OR social* OR arous*) NOT (cancer OR tumor OR stroke OR hematoma OR
hemorrhage OR aneurysm OR encephalitis OR infection)
```

**b** **Psychiatric Neuroimaging Corpus**

```
((positron emission tomography) OR “fmri”) AND (psychiat* OR “mental disorder” OR “mental
illness” OR “mental health” OR “neurodevelopmental disorder” OR “intellectual disability” OR
“communication disorder” OR autism OR adhd OR “learning disorder” OR “motor disorder” OR
psychotic OR psychosis OR delusion* OR schizophren* OR schizoaffect* OR catatoni* OR bipolar OR
“mood disorder” OR cyclothym* OR “mood dysregulation” OR “depressive disorder” OR depression OR
mdd OR “dysphoric disorder” OR anxiety OR anxious OR mutism OR phobia OR agoraphobia OR panic
OR ocd OR “obsessive-compulsive” OR “dysmorphic disorder” OR hoarding OR trichotillomania OR
skin-picking OR “reactive attachment disorder” OR “disinhibited social engagement” OR ptsd OR
“acute stress disorder” OR “adjustment disorder” OR “dissociative identity” OR “dissociative
amnesia” OR depersonalization OR derealization OR “dissociative disorder” OR ”somatic symptom”
OR “illness anxiety” OR “conversion disorder” OR “functional neurological symptom disorder” OR
“factitious disorder” OR “eating disorder” OR pica OR “rumination disorder” OR “restrictive
food” OR anorexia OR bulimia OR “binge eating” OR enuresis OR encopresis OR insomnia OR
hypersomnolence OR narcolepsy OR “sleep apnea” OR “sleep hypoventilation” OR “circadian rhythm
disorder” OR “sleep-wake disorder” OR “sleep arousal disorder” OR nightmare OR “sleep behavior
disorder” OR “restless legs syndrome” OR “sleep disorder” OR (“sexual dysfunction” AND brain)
OR (“delayed ejaculation“ AND brain) OR (“erectile dysfunction“ AND brain) OR (“orgasmic
disorder” AND brain) OR (“sexual interest” AND brain) OR (“sexual arousal” AND brain) OR
(“penetration disorder” AND brain) OR “sexual desire disorder” OR (“premature ejaculation” AND
brain) OR “gender dysphoria” OR “oppositional defiant disorder” OR “intermittent explosive
disorder” OR “conduct disorder” OR pyromania OR kleptomania OR “disruptive disorder” OR
“impulse-control disorder” OR “substance use disorder” OR “substance abuse” OR addiction OR
“alcohol use disorder” OR “alcohol intoxication” OR “alcohol withdrawal” OR “caffeine
intoxication” OR “caffeine withdrawal” OR “cannabis use disorder” OR “cannabis intoxication“ OR
“cannabis withdrawal” OR “phencyclidine use disorder” OR “hallucinogen use disorder” OR
“phencyclidine intoxication” OR “hallucinogen intoxication” OR “inherent use disorder” OR
“inhalant intoxication” OR “opioid use disorder” OR “opioid intoxication” OR “opioid
withdrawal” OR “sedative use disorder” OR “sedative intoxication” OR “sedative withdrawal” OR
“hypnotic use disorder” OR “hypnotic intoxication” OR “hypnotic withdrawal” OR “anxiolytic use
disorder” OR “anxiolytic intoxication” OR “anxiolytic withdrawal“ OR “stimulant use disorder“
OR “stimulant intoxication” OR “stimulant withdrawal” OR “tobacco use disorder” OR “tobacco
withdrawal“ OR delirium OR “neurocognitive disorder” OR “personality disorder“ OR “paranoid
personality” OR “schizoid personality” OR “schizotypal personality” OR “antisocial personality”
OR “borderline personality” OR “histrionic personality” OR “narcissistic personality“ OR
“avoidant personality” OR “dependent personality” OR “obsessive-compulsive personality” OR
paraphilia OR voyeurism OR exhibitionism OR frotteurism OR masochism OR sadism OR pedophilia OR
fetishism OR transvestism) NOT (cancer OR tumor OR stroke OR hematoma OR hemorrhage OR aneurysm
OR encephalitis OR infection)
```

**Extended Data Fig. 3 | Search queries for NLP corpora.** PubMed queries used to retrieve additional articles for **a,** the general neuroimaging corpus (see Extended Data Fig. 1b) and **b,** the psychiatric neuroimaging corpus (see Extended Data Fig. 1c). In addition to the above search criteria, studies were required to use human subjects and to be formatted as journal articles. Full text articles curated by these searches were added to the corpus of articles with coordinate data (see Extended Data Fig. 1a).

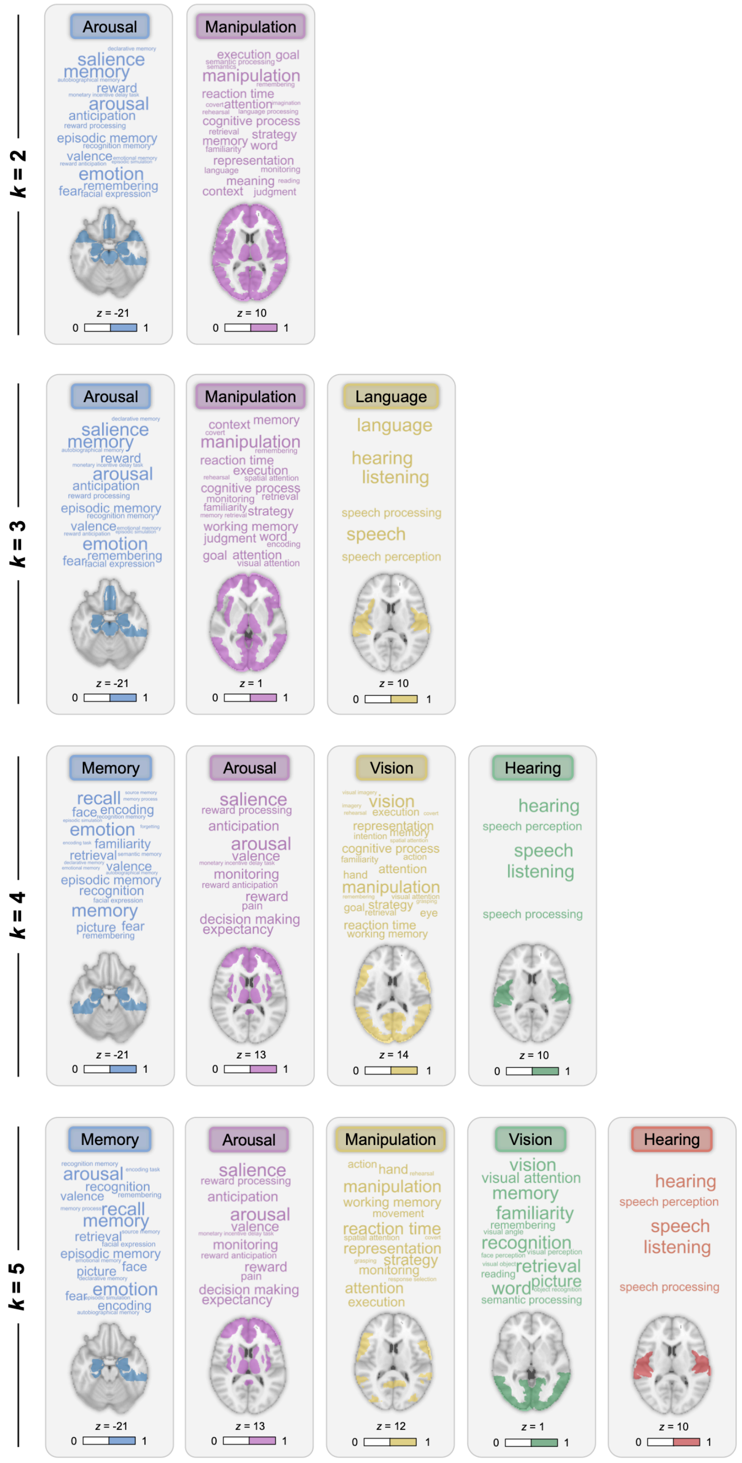


**Extended Data Fig. 4 | Data-driven solutions for *k*=2 to 5 using logistic regression classifiers to optimize term lists.** Domains generated at lower values of *k* than selected through the optimization procedure detailed in Fig. 1a.

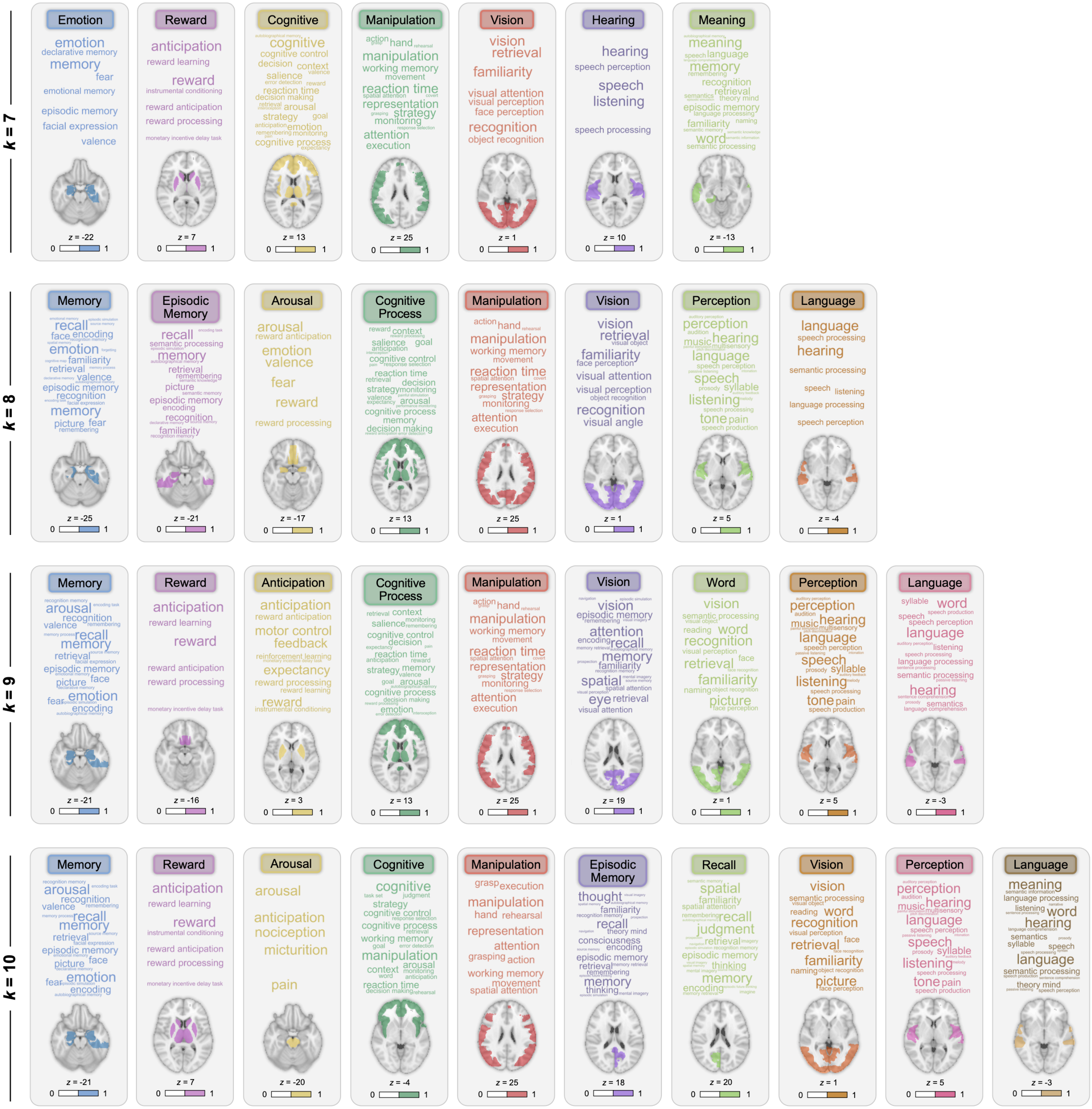


**Extended Data Fig. 5 | Data-driven solutions for *k*=7 to 10 using logistic regression classifiers to optimize term lists.** Domains generated at higher values of *k* than selected through the optimization procedure detailed in Fig. 1a.

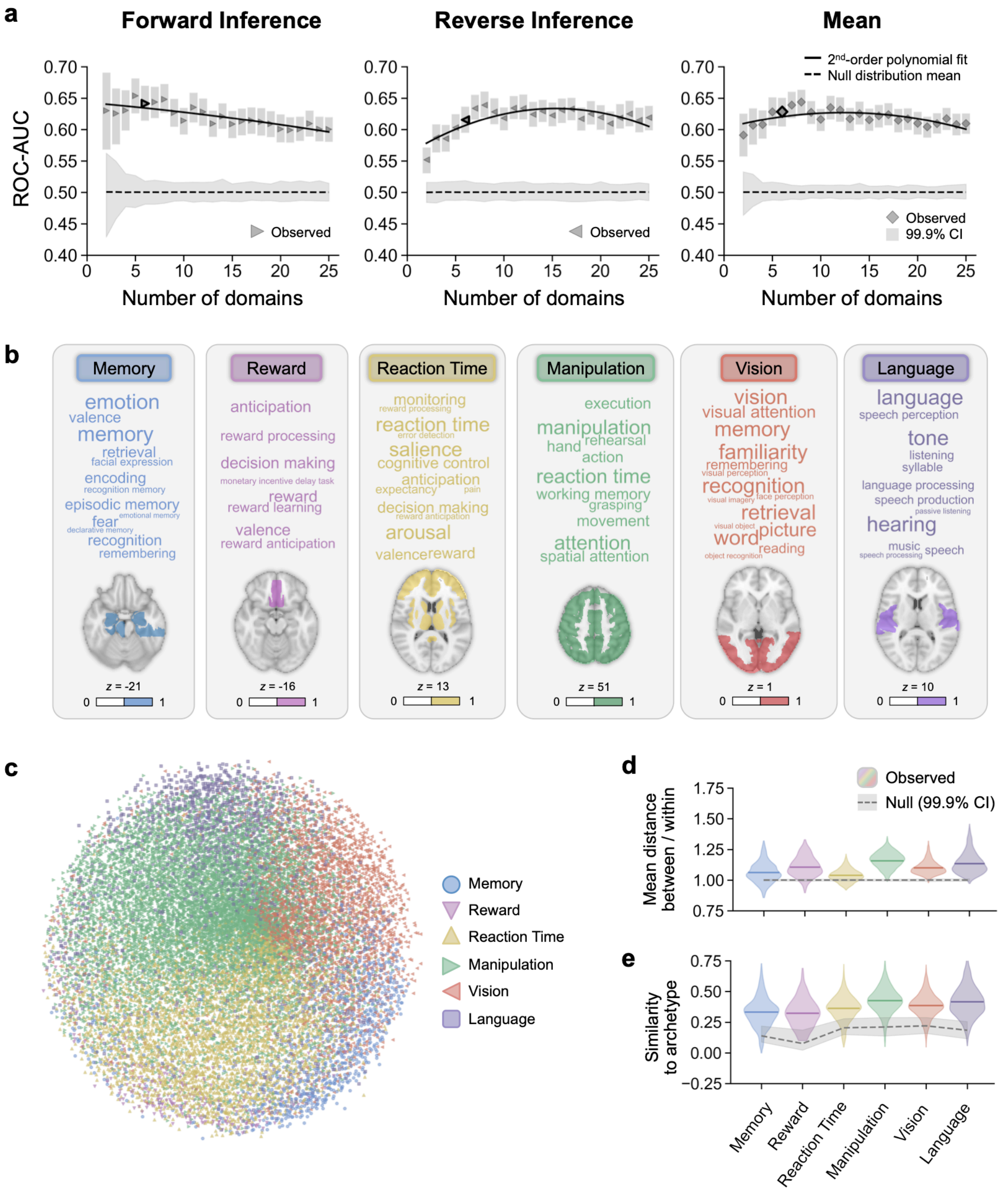


**Extended Data Fig. 6 | Neural network approach to data-driven ontology generation.** The procedure detailed in Fig. 1a was repeated using neural network classifiers in place of logistic regression. All neural network classifiers were comprised of 8 fully connected layers and were fit with learning rate = 0.001, weight decay = 0.001, neurons per layer = 100, dropout probability = 0.1 (applied to the last 3 layers), and batch size = 1,024. In Step 3 of the process, classifiers predicted term and structure occurrences within domains and were trained over 100 epochs. In Step 4, classifiers predicted domain word list and circuit occurrences and were trained over 500 epochs. **a,** Validation set ROC-AUC for logistic regression classifiers used to select the optimal number of domains. Performance is plotted for forward inference classifiers (right-pointing triangles), reverse inference classifiers (left-pointing triangles), and their average (diamonds). Shaded areas around markers represent 99.9% confidence intervals computed by resampling validation set articles with replacement over 1,000 iterations. The dashed line represents the mean of null distributions generated by shuffling true labels for validation set articles over 1,000 iterations, and the surrounding shaded area is the 95% confidence interval. **b,** Data-driven solution for 6 domains. Word size is scaled to frequency in the corpus of 18,155 articles with activation coordinate data. The number of words per domain was selected in Step 3 using logistic regression performance in the validation set. Brain maps show structures included in each circuit as a result of clustering by PMI-weighted co-occurrences with function terms, which were not influenced by the downstream neural network approach. **c,** Article partitioning based on maximal similarity to terms and structures in domain archetypes plotted with multidimensional scaling. **d,** Modularity of the article partitioning was assessed by comparing the mean Dice distance of function and structure occurrences of articles between domains versus within domains. Observed values are colored by domain; null distributions in gray were computed by shuffling distance values across article partitions. **e,** Generalizability was assessed by computing the Dice similarity of each domain's "archetype" vector of function terms and brain structures with the terms and structure occurring in each article of the domain's partition. Observed values are colored by domain; null distributions in gray were computed by shuffling terms and structures in each archetype.

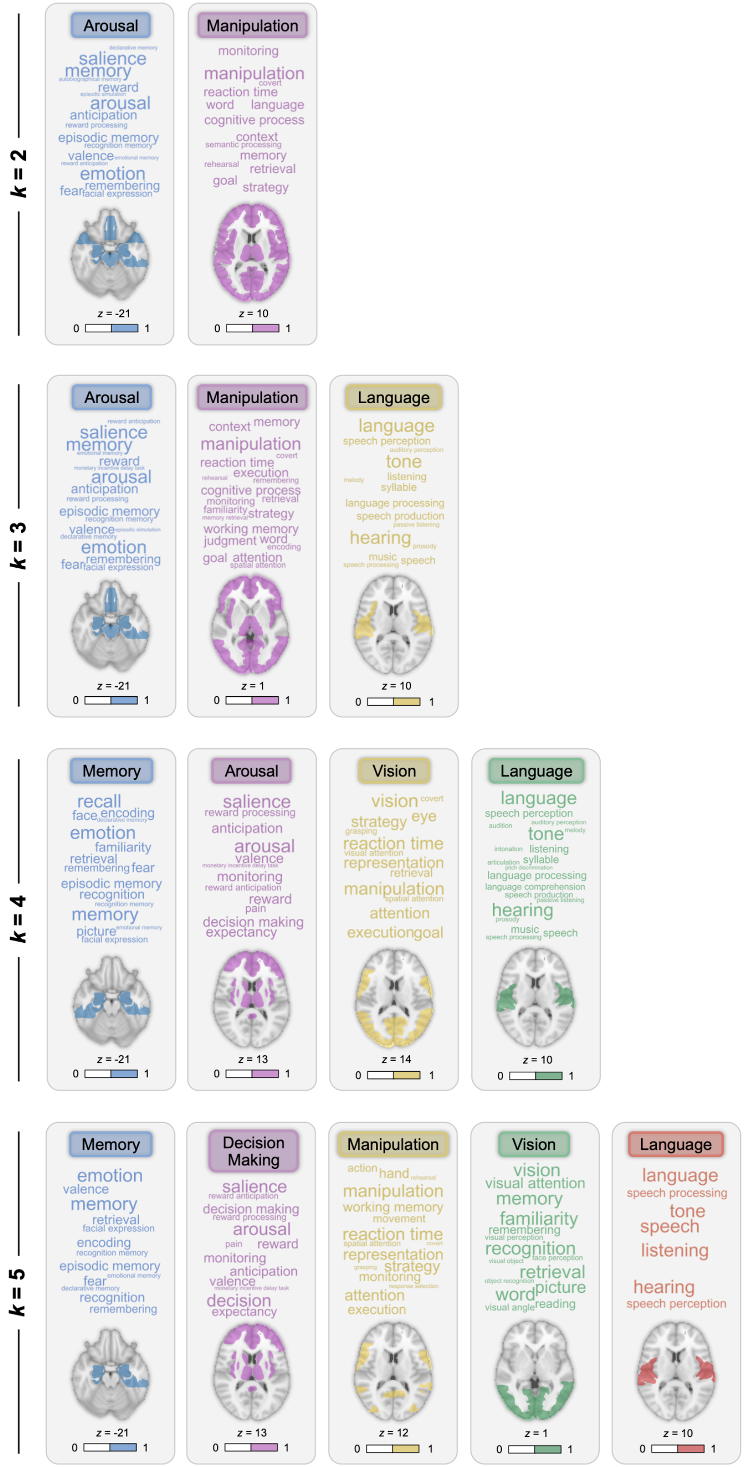


**Extended Data Fig. 7 | Data-driven solutions for *k*=2 to 5 using neural network classifiers for optimizing term lists.** Domains generated at lower values of *k* than selected based on ROC-AUC results shown in Fig. 6a.

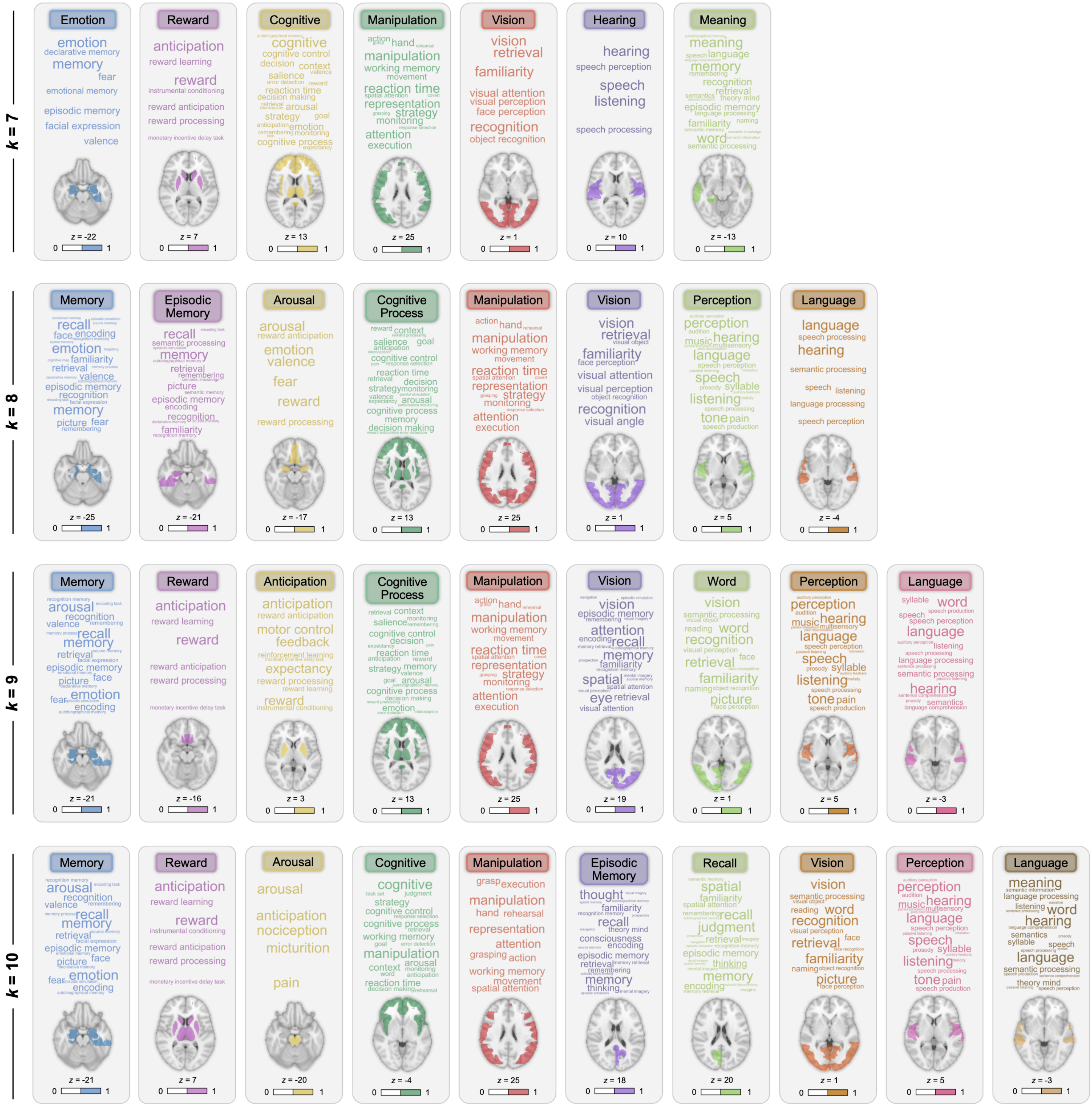


**Extended Data Fig. 8 | Data-driven solutions for $k$=7 to 10 using neural network classifiers for optimizing term lists.** Domains generated at lower values of $k$ than selected based on ROC-AUC results shown in Fig. 6a.

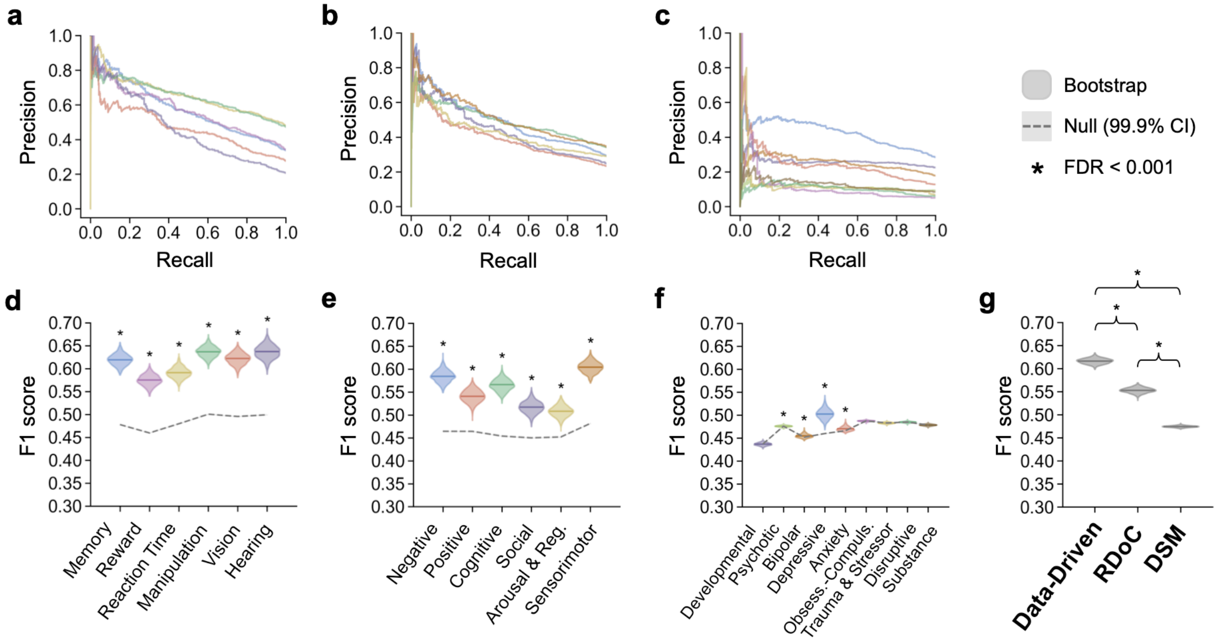


**Extended Data Fig. 9 | Additional evaluation metrics for reverse inference classification with logistic regression.** Precision-recall curves are shown for the test set performance of logistic regression classifiers with mental function features defined by **a,** the data-driven ontology, **b,** RDoC, and **c,** the DSM. **d-f,** Bootstrap distributions of F1 score (colored) were computed by resampling articles in the test set with replacement over 1,000 iterations. Observed values in the test set are shown with solid lines. Null distributions (gray) were computed by shuffling true labels for term list scores over 1,000 iterations; the 99.9% confidence interval is shaded, and distribution means are shown with dashed lines. The mean of each bootstrap distribution was assessed for a difference in mean from its corresponding null distribution (* FDR < 0.001). **g,** F1 scores across the domains in each framework. Solid lines denote means of the bootstrap distributions macro-averaged across classifiers. Differences in bootstrap means were assessed for each framework pair (* FDR < 0.001).

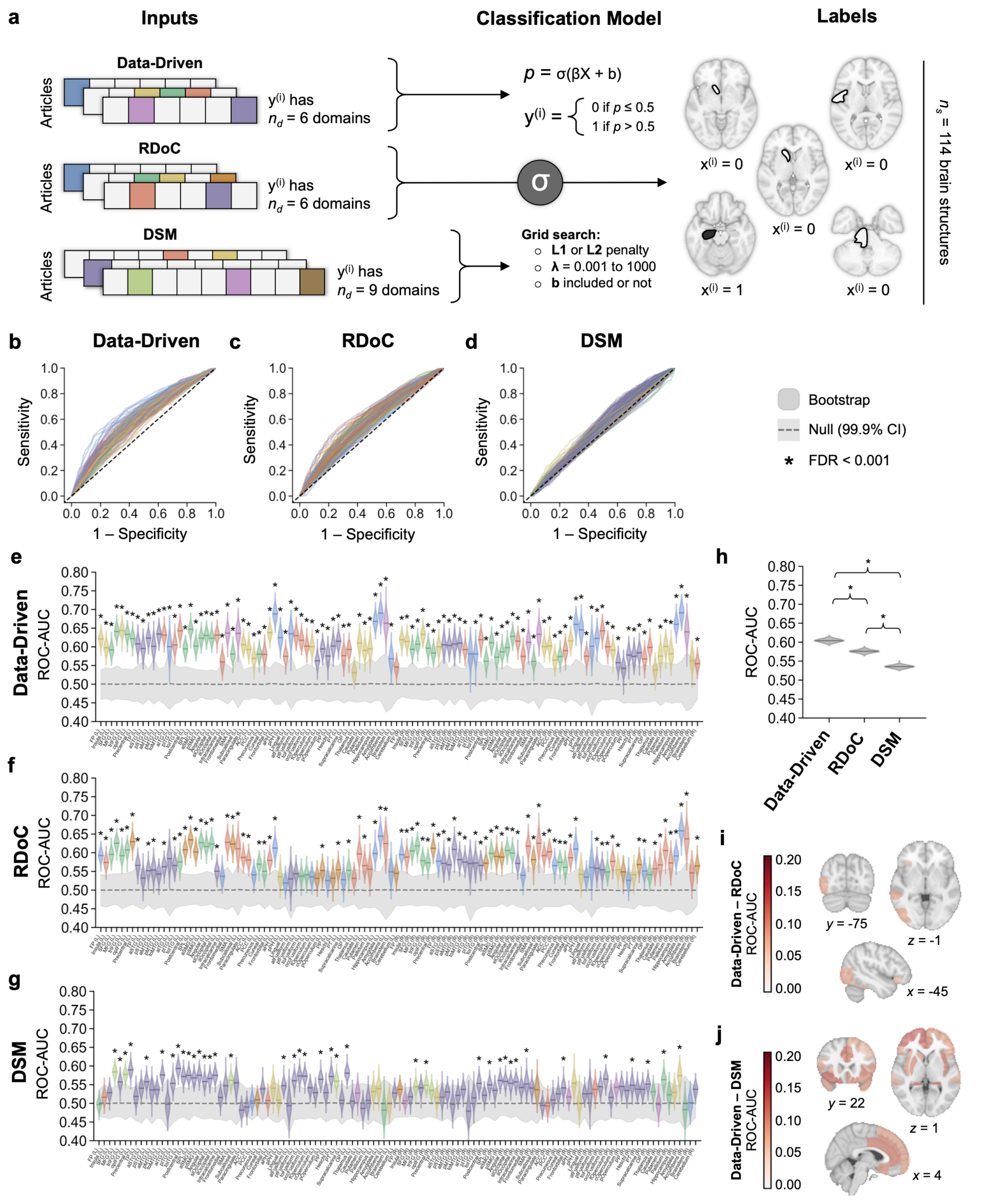


**Extended Data Fig. 10 | Forward inference classification with logistic regression. a,** Logistic regression classifiers were trained to predict whether coordinates were reported in each of 114 brain structures based on the occurrences of words for mental functions in neuroimaging article full texts. Plots are colored by the domain assignment for structures in the data-driven framework, and by the domain with the highest PPMI for the structure in RDoC and DSM frameworks. Classifier features included word occurrences thresholded by mean frequency across the corpus, then the mean frequency of words in each domain. Activation coordinate data were mapped to anatomically defined structures the brain[2] and cerebellum.[3] Training was performed in 70% of articles (*n* = 12,708), hyperparameters were tuned on a validation set containing 20% of articles (*n* = 3,631), then classifiers were evaluated in a test set containing 10% of articles (*n* = 1,816). ROC curves are shown for the test set performance of classifiers with mental function features defined by **b,** the data-driven ontology, **c,** RDoC, and **d,** the DSM. **e-g,** Bootstrap distributions of ROC-AUC (colored) were computed by resampling articles in the test set over 1,000 iterations. Observed values in the test set are shown with solid lines. Null distributions (gray) were computed by shuffling true labels over 1,000 iterations; the 99.9% confidence interval is shaded, and distribution means are shown with dashed lines. Bootstrap distributions were assessed for overlap with null distributions (* FDR < 0.001). **h,** ROC-AUC across the domains in each framework. Solid lines denote means of the bootstrap distributions macro-averaged across classifiers. Differences in bootstrap means were assessed for each framework pair (* FDR < 0.001). **I-j,** Difference in ROC-AUC between the data-driven and expert determined frameworks. Maps were thresholded to show differences with FDR < 0.001 based on permutation testing.

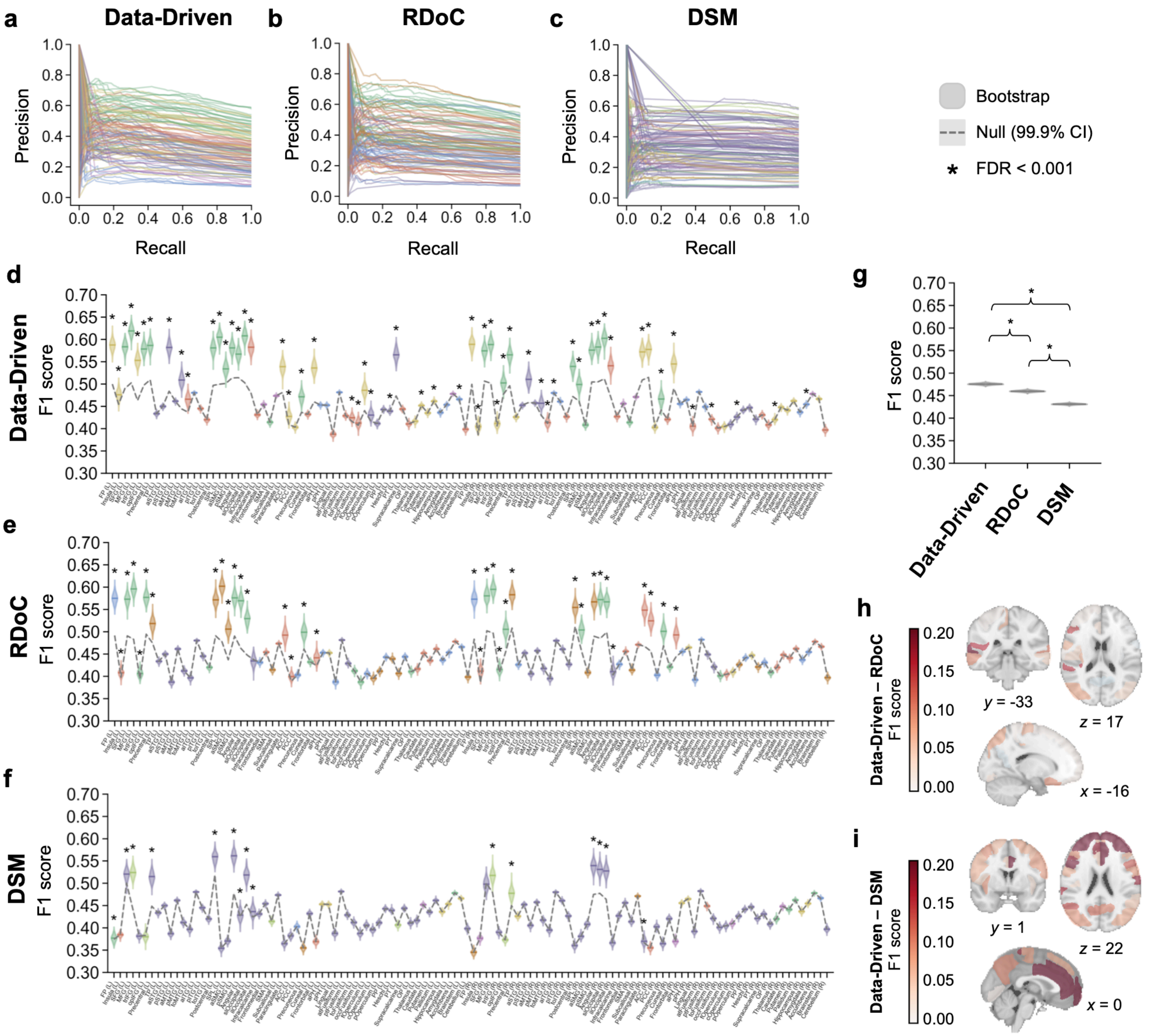


**Extended Data Fig. 11 | Additional evaluation metrics for forward inference classification with logistic regression.** Precision-recall curves are shown for the test set performance of logistic regression classifiers predicting mental function term lists defined by **a,** the data-driven ontology, **b,** RDoC, and **c,** the DSM. **d-f,** Bootstrap distributions of F1 score (colored) were computed by resampling articles in the test set with replacement over 1,000 iterations. Observed values in the test set are shown with solid lines. Null distributions (gray) were computed by shuffling true labels for term list scores over 1,000 iterations; the 99.9% confidence interval is shaded, and distribution means are shown with dashed lines. The mean of each bootstrap distribution was assessed for a difference in mean from its corresponding null distribution (* FDR < 0.001). **g,** F1 scores across the domains in each framework. Solid lines denote means of the bootstrap distributions macro-averaged across classifiers. Differences in bootstrap means were assessed for each framework pair (* FDR < 0.001). **I-j,** Difference in F1 scores between the data-driven and expert determined frameworks. Maps were thresholded to show differences with FDR < 0.001 based on permutation testing.

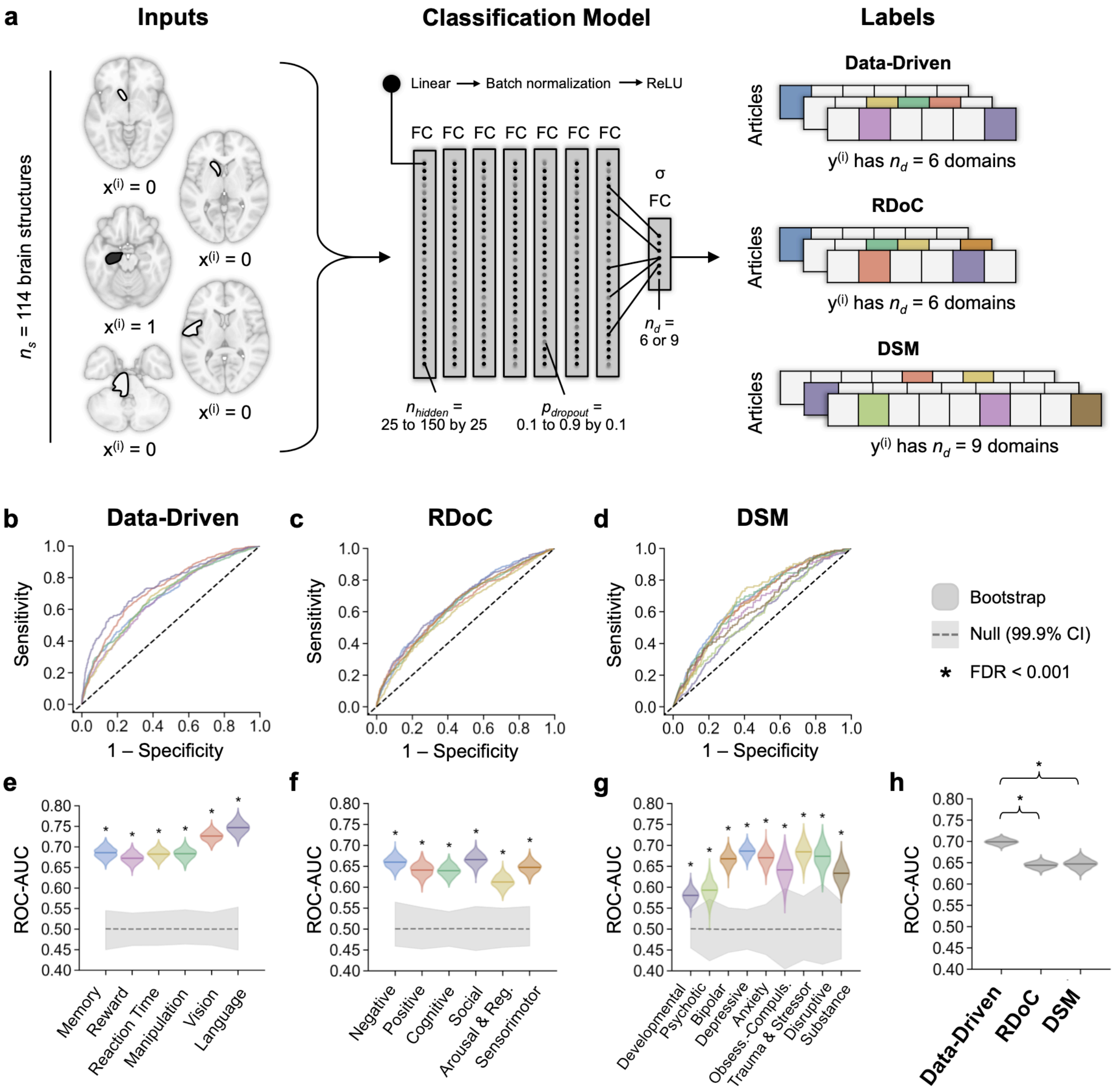


**Extended Data Fig. 12 | Reverse inference classification with neural networks.** Neural network classifiers were trained to perform reverse inference with the same features and labels as described in Fig. 4. Classification models comprised 8 fully connected (FC), all with ReLU activation functions except the output layer which was activated by a sigmoid. The optimal learning rate, weight decay, number of neurons per layer, and dropout probability were determined for each framework through a randomized grid search. ROC curves are shown for the test set performance of classifiers with mental function features defined by **b,** the data-driven ontology, **c,** RDoC, and **d,** the DSM. **e-g,** Bootstrap distributions of ROC-AUC (colored) were computed by resampling articles in the test set with replacement over 1,000 iterations. Observed values in the test set are shown with solid lines. Null distributions (gray) were computed by shuffling true labels for term list scores over 1,000 iterations; the 99.9% confidence interval is shaded, and distribution means are shown with dashed lines. The mean of each bootstrap distribution was assessed for a difference in mean from its corresponding null distribution (* FDR < 0.001). **h,** ROC-AUC across the domains in each framework. Solid lines denote means of the bootstrap distributions macro-averaged across classifiers. Differences in bootstrap means were assessed for each framework pair (* FDR < 0.001).

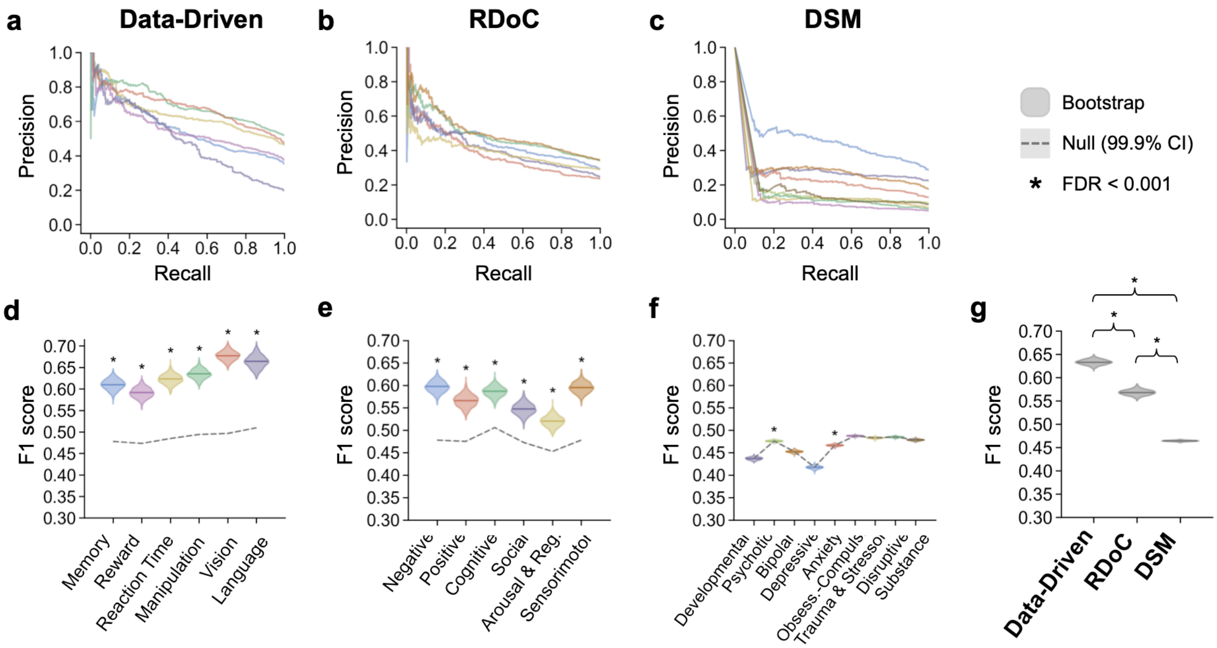


**Extended Data Fig. 13 | Additional evaluation metrics for reverse inference classification with neural networks.** Precision-recall curves are shown for the test set performance of neural network classifiers with mental function features defined by **a,** the data-driven ontology, **b,** RDoC, and **c,** the DSM. **d-f,** Bootstrap distributions of F1 score (colored) were computed by resampling articles in the test set with replacement over 1,000 iterations. Observed values in the test set are shown with solid lines. Null distributions (gray) were computed by shuffling true labels for term list scores over 1,000 iterations; the 99.9% confidence interval is shaded, and distribution means are shown with dashed lines. The mean of each bootstrap distribution was assessed for a difference in mean from its corresponding null distribution (* FDR < 0.001). **g,** F1 scores across the domains in each framework. Solid lines denote means of the bootstrap distributions macro-averaged across classifiers. Differences in bootstrap means were assessed for each framework pair (* FDR < 0.001).

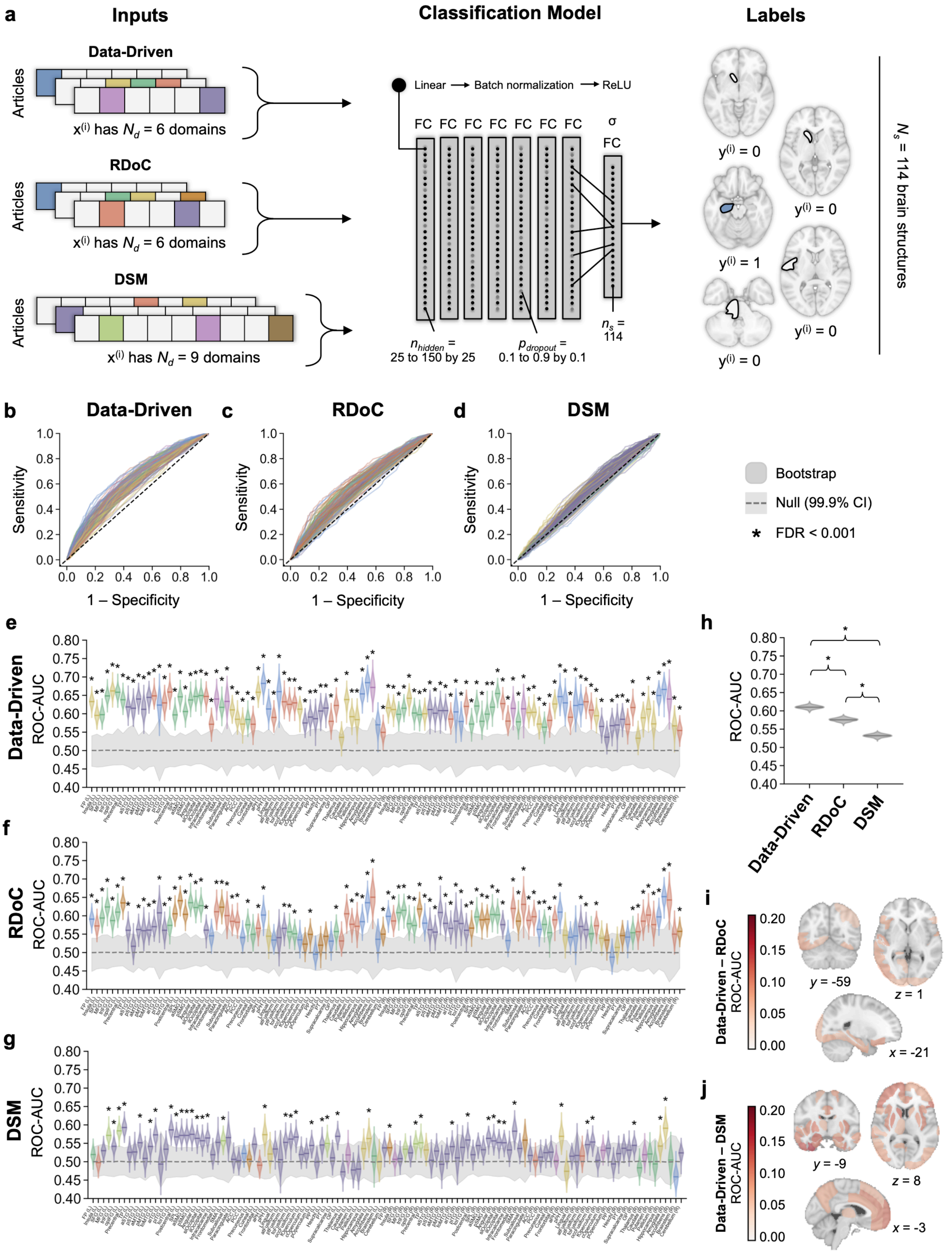
a
Inputs
Classification Model
Labels
Data-Driven
Articles
x(i) has Nd = 6 domains
RDoC
Articles
x(i) has Nd = 6 domains
DSM
Articles
x(i) has Nd = 9 domains
Linear → Batch normalization → ReLU
FC FC FC FC FC FC FC
σ
FC
nhidden = 25 to 150 by 25
pdropout = 0.1 to 0.9 by 0.1
ns = 114
y(i) = 0
y(i) = 0
y(i) = 1
y(i) = 0
y(i) = 0
Ns = 114 brain structures
b
Data-Driven
c
RDoC
d
DSM
Sensitivity
1 – Specificity
Bootstrap
Null (99.9% CI)
* FDR < 0.001
e
Data-Driven
ROC-AUC
f
RDoC
ROC-AUC
g
DSM
ROC-AUC
h
ROC-AUC
Data-Driven
RDoC
DSM
i
Data-Driven – RDoC
ROC-AUC
y = -59
z = 1
x = -21
j
Data-Driven – DSM
ROC-AUC
y = -9
z = 8
x = -3

**Extended Data Fig. 14 | Forward inference classification with neural networks. a,** Neural network classifiers were trained to perform forward inference with the same features and labels as described in Extended Data Fig. 8. Forward inference classifiers were optimized over a grid search with the same hyperparameter values as reverse inference classifiers in Extended Data Fig. 10. ROC curves are shown for the test set performance of classifiers with mental function features defined by **b,** the data-driven ontology, **c,** RDoC, and **d,** the DSM. **e-g,** Bootstrap distributions of ROC-AUC (colored) were computed by resampling articles in the test set over 1,000 iterations. Observed values in the test set are shown with solid lines. Null distributions (gray) were computed by shuffling true labels over 1,000 iterations; the 99.9% confidence interval is shaded, and distribution means are shown with dashed lines. Bootstrap distributions were assessed for overlap with null distributions (* FDR < 0.001). **h,** ROC-AUC across the domains in each framework. Solid lines denote means of the bootstrap distributions macro-averaged across classifiers. Differences in bootstrap means were assessed for each framework pair (* FDR < 0.001). **I-j,** Difference in ROC-AUC between the data-driven and expert determined frameworks. Maps were thresholded to show differences with FDR < 0.001 based on permutation testing.

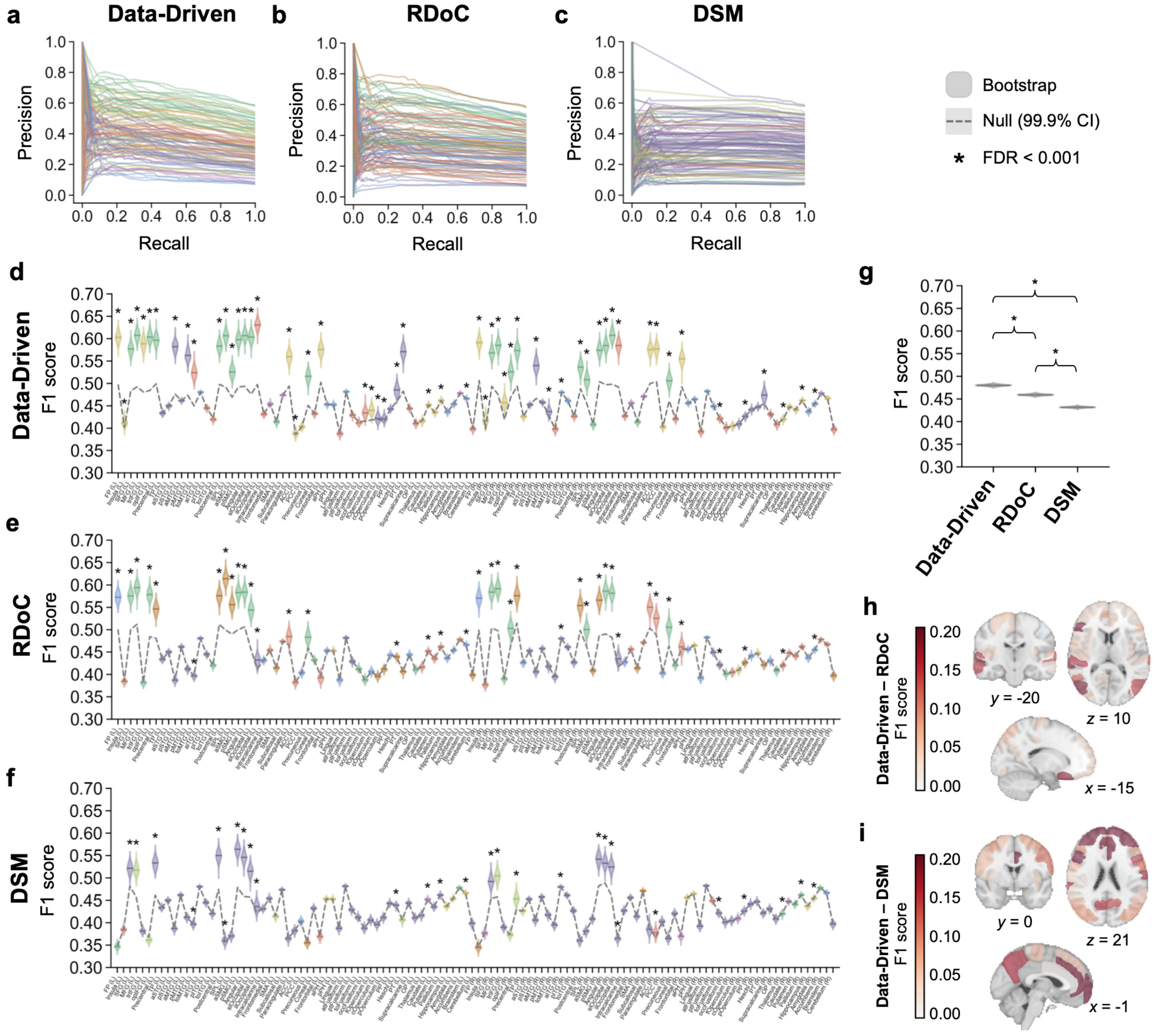


**Extended Data Fig. 15 | Additional evaluation metrics for forward inference classification with neural networks.** Precision-recall curves are shown for the test set performance of neural network classifiers predicting mental function term lists defined by **a,** the data-driven ontology, **b,** RDoC, and **c,** the DSM. **d-f,** Bootstrap distributions of F1 score (colored) were computed by resampling articles in the test set with replacement over 1,000 iterations. Observed values in the test set are shown with solid lines. Null distributions (gray) were computed by shuffling true labels for term list scores over 1,000 iterations; the 99.9% confidence interval is shaded, and distribution means are shown with dashed lines. The mean of each bootstrap distribution was assessed for a difference in mean from its corresponding null distribution (* FDR < 0.001). **g,** F1 scores across the domains in each framework. Solid lines denote means of the bootstrap distributions macro-averaged across classifiers. Differences in bootstrap means were assessed for each framework pair (* FDR < 0.001). **I-j,** Difference in F1 scores between the data-driven and expert determined frameworks. Maps were thresholded to show differences with FDR < 0.001 based on permutation testing.

### Data-Driven

| | ROC-AUC | | | F1 | | |
|---|---|---|---|---|---|---|
| | LR | NN | 99.9% CI | LR | NN | 99.9% CI |
| Memory | 0.71 | 0.69 | -0.03 to 0.09 | 0.62 | 0.61 | -0.04 to 0.06 |
| Reward | 0.71 | 0.67 | -0.02 to 0.10 | 0.58 | 0.59 | -0.06 to 0.04 |
| Reaction Time | 0.68 | 0.68 | -0.06 to 0.05 | 0.59 | 0.62 | -0.08 to 0.02 |
| Manipulation | 0.68 | 0.68 | -0.06 to 0.06 | 0.64 | 0.64 | -0.04 to 0.05 |
| Vision | 0.70 | 0.73 | -0.08 to 0.04 | 0.62 | 0.68 | -0.11 to 0.01 |
| Hearing/Language | 0.72 | 0.75 | -0.09 to 0.03 | 0.64 | 0.66 | -0.09 to 0.05 |

### RDoC

| | ROC-AUC | | | F1 | | |
|---|---|---|---|---|---|---|
| | LR | NN | 99.9% CI | LR | NN | 99.9% CI |
| Negative Valence | 0.69 | 0.66 | -0.03 to 0.09 | 0.58 | 0.60 | -0.07 to 0.04 |
| Positive Valence | 0.67 | 0.64 | -0.03 to 0.11 | 0.54 | 0.57 | -0.08 to 0.04 |
| Cognitive Systems | 0.66 | 0.64 | -0.03 to 0.08 | 0.57 | 0.59 | -0.07 to 0.03 |
| Social Processes | 0.68 | 0.67 | -0.04 to 0.09 | 0.52 | 0.55 | -0.08 to 0.03 |
| Arousal & Regulation | 0.63 | 0.61 | -0.04 to 0.10 | 0.51 | 0.52 | -0.06 to 0.05 |
| Sensorimotor Systems | 0.65 | 0.65 | -0.05 to 0.07 | 0.60 | 0.60 | -0.04 to 0.07 |

### DSM

| | ROC-AUC | | | F1 | | |
|---|---|---|---|---|---|---|
| | LR | NN | 99.9% CI | LR | NN | 99.9% CI |
| Developmental | 0.56 | 0.58 | -0.07 to 0.06 | 0.44 | 0.44 | -0.02 to 0.01 |
| Psychotic | 0.56 | 0.59 | -0.14 to 0.08 | 0.48 | 0.48 | -0.01 to 0.01 |
| Bipolar | 0.65 | 0.67 | -0.09 to 0.09 | 0.45 | 0.45 | -0.01 to 0.02 |
| Depressive | 0.67 | 0.69 | -0.07 to 0.05 | 0.50 | 0.42 | 0.05 to 0.13 |
| Anxiety | 0.68 | 0.67 | -0.07 to 0.09 | 0.47 | 0.47 | -0.01 to 0.03 |
| Obsessive-Compulsive | 0.64 | 0.64 | -0.12 to 0.13 | 0.49 | 0.49 | -0.01 to 0.01 |
| Trauma & Stressor | 0.67 | 0.68 | -0.10 to 0.09 | 0.48 | 0.48 | -0.01 to 0.01 |
| Disruptive | 0.66 | 0.67 | -0.14 to 0.14 | 0.48 | 0.48 | -0.01 to 0.01 |
| Substance | 0.62 | 0.63 | -0.10 to 0.11 | 0.48 | 0.48 | -0.01 to 0.01 |

**Extended Data Fig. 16 | Reverse inference results across classification architectures.** Observed test set ROC-AUC and F1 scores for logistic regression (LR) and neural network (NN) classifiers. Each 99.9% confidence interval (CI) for the difference between LR and NN classifiers was computed by bootstrap resampling of articles in the test set over 1,000 iterations. Classifiers for which the 99.9% CI did not include zero are highlighted in yellow.

| | Data-Driven | | | | | | RDoC | | | | | | DSM | | | | | |
|---|---|---|---|---|---|---|---|---|---|---|---|---|---|---|---|---|---|---|
| | ROC-AUC | | | F1 | | | ROC-AUC | | | F1 | | | ROC-AUC | | | F1 | | |
| | LR | NN | 99.9% CI | LR | NN | 99.9% CI | LR | NN | 99.9% CI | LR | NN | 99.9% CI | LR | NN | 99.9% CI | LR | NN | 99.9% CI |
| FP (L) | 0.62 | 0.63 | -0.07 to 0.04 | 0.59 | 0.60 | -0.07 to 0.03 | 0.59 | 0.59 | -0.06 to 0.07 | 0.58 | 0.57 | -0.04 to 0.05 | 0.50 | 0.52 | -0.08 to 0.03 | 0.38 | 0.35 | 0.00 to 0.06 |
| Insula (L) | 0.60 | 0.60 | -0.06 to 0.07 | 0.48 | 0.41 | 0.02 to 0.11 | 0.57 | 0.57 | -0.06 to 0.07 | 0.41 | 0.38 | -0.01 to 0.05 | 0.52 | 0.50 | -0.05 to 0.10 | 0.38 | 0.38 | -0.02 to 0.02 |
| SFG (L) | 0.59 | 0.60 | -0.07 to 0.06 | 0.58 | 0.58 | -0.04 to 0.06 | 0.60 | 0.59 | -0.06 to 0.07 | 0.57 | 0.58 | -0.05 to 0.06 | 0.53 | 0.53 | -0.05 to 0.05 | 0.52 | 0.52 | -0.05 to 0.05 |
| MFG (L) | 0.64 | 0.65 | -0.06 to 0.05 | 0.62 | 0.61 | -0.04 to 0.06 | 0.63 | 0.62 | -0.05 to 0.06 | 0.60 | 0.59 | -0.05 to 0.07 | 0.58 | 0.57 | -0.04 to 0.07 | 0.52 | 0.52 | -0.04 to 0.06 |
| trIFG (L) | 0.64 | 0.66 | -0.07 to 0.04 | 0.55 | 0.59 | -0.08 to 0.02 | 0.59 | 0.59 | -0.04 to 0.07 | 0.41 | 0.38 | 0.00 to 0.05 | 0.56 | 0.54 | -0.04 to 0.08 | 0.38 | 0.38 | -0.02 to 0.02 |
| opIFG (L) | 0.63 | 0.66 | -0.07 to 0.03 | 0.58 | 0.60 | -0.08 to 0.03 | 0.61 | 0.61 | -0.06 to 0.06 | 0.58 | 0.58 | -0.05 to 0.05 | 0.58 | 0.58 | -0.06 to 0.06 | 0.38 | 0.36 | -0.01 to 0.05 |
| Precentral (L) | 0.62 | 0.64 | -0.08 to 0.05 | 0.59 | 0.60 | -0.07 to 0.05 | 0.63 | 0.64 | -0.07 to 0.06 | 0.52 | 0.55 | -0.08 to 0.02 | 0.59 | 0.59 | -0.06 to 0.05 | 0.51 | 0.53 | -0.07 to 0.03 |
| TP (L) | 0.61 | 0.62 | -0.08 to 0.07 | 0.43 | 0.43 | -0.01 to 0.02 | 0.57 | 0.56 | -0.06 to 0.07 | 0.43 | 0.43 | -0.01 to 0.02 | 0.52 | 0.53 | -0.07 to 0.06 | 0.43 | 0.43 | -0.01 to 0.02 |
| aSTG (L) | 0.60 | 0.61 | -0.09 to 0.06 | 0.45 | 0.45 | -0.01 to 0.01 | 0.53 | 0.52 | -0.06 to 0.09 | 0.45 | 0.45 | -0.01 to 0.01 | 0.54 | 0.53 | -0.05 to 0.09 | 0.45 | 0.45 | -0.01 to 0.01 |
| pSTG (L) | 0.62 | 0.64 | -0.07 to 0.05 | 0.58 | 0.58 | -0.05 to 0.06 | 0.55 | 0.56 | -0.07 to 0.05 | 0.39 | 0.39 | -0.02 to 0.02 | 0.56 | 0.55 | -0.05 to 0.06 | 0.39 | 0.39 | -0.02 to 0.02 |
| aMTG (L) | 0.60 | 0.61 | -0.09 to 0.09 | 0.46 | 0.46 | -0.01 to 0.01 | 0.55 | 0.56 | -0.08 to 0.07 | 0.46 | 0.46 | -0.01 to 0.01 | 0.54 | 0.51 | -0.05 to 0.09 | 0.46 | 0.46 | -0.01 to 0.01 |
| pMTG (L) | 0.63 | 0.64 | -0.07 to 0.06 | 0.51 | 0.56 | -0.10 to 0.00 | 0.54 | 0.56 | -0.08 to 0.04 | 0.41 | 0.41 | -0.02 to 0.02 | 0.54 | 0.54 | -0.06 to 0.07 | 0.41 | 0.41 | -0.02 to 0.02 |
| toMTG (L) | 0.63 | 0.65 | -0.09 to 0.05 | 0.47 | 0.52 | -0.11 to -0.01 | 0.56 | 0.57 | -0.06 to 0.05 | 0.40 | 0.40 | -0.02 to 0.02 | 0.58 | 0.57 | -0.05 to 0.08 | 0.40 | 0.40 | -0.02 to 0.02 |
| aITG (L) | 0.60 | 0.62 | -0.13 to 0.10 | 0.48 | 0.48 | -0.01 to 0.01 | 0.58 | 0.61 | -0.14 to 0.09 | 0.48 | 0.48 | -0.01 to 0.01 | 0.50 | 0.49 | -0.09 to 0.12 | 0.48 | 0.48 | -0.01 to 0.01 |
| pITG (L) | 0.61 | 0.63 | -0.09 to 0.06 | 0.44 | 0.44 | -0.01 to 0.01 | 0.57 | 0.56 | -0.06 to 0.09 | 0.44 | 0.44 | -0.01 to 0.01 | 0.55 | 0.53 | -0.04 to 0.08 | 0.44 | 0.44 | -0.01 to 0.01 |
| toITG (L) | 0.64 | 0.66 | -0.08 to 0.05 | 0.42 | 0.42 | -0.01 to 0.02 | 0.58 | 0.57 | -0.05 to 0.09 | 0.42 | 0.42 | -0.01 to 0.02 | 0.59 | 0.59 | -0.04 to 0.06 | 0.42 | 0.42 | -0.01 to 0.02 |
| Postcentral (L) | 0.59 | 0.60 | -0.06 to 0.05 | 0.58 | 0.58 | -0.05 to 0.05 | 0.61 | 0.61 | -0.06 to 0.06 | 0.57 | 0.58 | -0.05 to 0.05 | 0.57 | 0.57 | -0.05 to 0.06 | 0.56 | 0.55 | -0.04 to 0.08 |
| SPL (L) | 0.65 | 0.65 | -0.05 to 0.08 | 0.60 | 0.61 | -0.04 to 0.06 | 0.63 | 0.64 | -0.06 to 0.06 | 0.60 | 0.61 | -0.06 to 0.03 | 0.58 | 0.57 | -0.05 to 0.06 | 0.35 | 0.36 | -0.03 to 0.02 |
| aSMG (L) | 0.60 | 0.61 | -0.06 to 0.05 | 0.53 | 0.53 | -0.04 to 0.06 | 0.60 | 0.60 | -0.06 to 0.06 | 0.51 | 0.56 | -0.10 to 0.01 | 0.57 | 0.57 | -0.05 to 0.05 | 0.37 | 0.37 | -0.02 to 0.02 |
| pSMG (L) | 0.62 | 0.64 | -0.07 to 0.04 | 0.58 | 0.60 | -0.06 to 0.04 | 0.63 | 0.64 | -0.07 to 0.06 | 0.58 | 0.58 | -0.05 to 0.06 | 0.58 | 0.57 | -0.05 to 0.06 | 0.56 | 0.56 | -0.05 to 0.05 |
| Angular (L) | 0.63 | 0.65 | -0.07 to 0.04 | 0.57 | 0.61 | -0.08 to 0.01 | 0.62 | 0.62 | -0.06 to 0.05 | 0.57 | 0.58 | -0.06 to 0.04 | 0.56 | 0.56 | -0.06 to 0.05 | 0.43 | 0.55 | -0.17 to -0.06 |
| slOccipital (L) | 0.62 | 0.65 | -0.09 to 0.02 | 0.61 | 0.60 | -0.04 to 0.06 | 0.62 | 0.63 | -0.06 to 0.06 | 0.53 | 0.54 | -0.07 to 0.05 | 0.57 | 0.56 | -0.04 to 0.06 | 0.52 | 0.52 | -0.04 to 0.07 |
| ilOccipital (L) | 0.63 | 0.65 | -0.06 to 0.05 | 0.58 | 0.63 | -0.09 to 0.02 | 0.55 | 0.56 | -0.07 to 0.05 | 0.43 | 0.43 | -0.04 to 0.05 | 0.57 | 0.56 | -0.04 to 0.06 | 0.44 | 0.44 | -0.04 to 0.05 |
| Intracalcarine (L) | 0.56 | 0.57 | -0.08 to 0.06 | 0.43 | 0.43 | -0.01 to 0.01 | 0.54 | 0.55 | -0.07 to 0.08 | 0.43 | 0.43 | -0.01 to 0.01 | 0.52 | 0.52 | -0.06 to 0.05 | 0.43 | 0.43 | -0.01 to 0.01 |
| Frontomedial (L) | 0.64 | 0.64 | -0.07 to 0.07 | 0.45 | 0.45 | -0.01 to 0.01 | 0.63 | 0.61 | -0.06 to 0.09 | 0.45 | 0.45 | -0.01 to 0.01 | 0.55 | 0.55 | -0.07 to 0.07 | 0.45 | 0.45 | -0.01 to 0.01 |
| SMA (L) | 0.58 | 0.62 | -0.10 to 0.02 | 0.41 | 0.41 | -0.02 to 0.02 | 0.62 | 0.62 | -0.06 to 0.07 | 0.41 | 0.41 | -0.02 to 0.02 | 0.56 | 0.56 | -0.05 to 0.07 | 0.41 | 0.41 | -0.02 to 0.02 |
| Subcallosal (L) | 0.64 | 0.63 | -0.08 to 0.10 | 0.47 | 0.47 | -0.01 to 0.01 | 0.61 | 0.60 | -0.07 to 0.11 | 0.47 | 0.47 | -0.01 to 0.01 | 0.55 | 0.57 | -0.09 to 0.08 | 0.47 | 0.47 | -0.01 to 0.01 |
| Paracingulate (L) | 0.61 | 0.61 | -0.06 to 0.06 | 0.54 | 0.56 | -0.07 to 0.03 | 0.59 | 0.59 | -0.06 to 0.08 | 0.49 | 0.48 | -0.04 to 0.06 | 0.48 | 0.51 | -0.08 to 0.03 | 0.36 | 0.36 | -0.02 to 0.02 |
| ACC (L) | 0.59 | 0.59 | -0.05 to 0.07 | 0.43 | 0.39 | 0.01 to 0.07 | 0.58 | 0.58 | -0.06 to 0.06 | 0.40 | 0.38 | -0.01 to 0.05 | 0.49 | 0.50 | -0.06 to 0.05 | 0.38 | 0.38 | -0.02 to 0.02 |
| PCC (L) | 0.57 | 0.57 | -0.06 to 0.07 | 0.40 | 0.40 | -0.02 to 0.02 | 0.54 | 0.54 | -0.06 to 0.06 | 0.40 | 0.40 | -0.02 to 0.02 | 0.51 | 0.52 | -0.08 to 0.04 | 0.40 | 0.40 | -0.02 to 0.02 |
| Precuneous (L) | 0.58 | 0.58 | -0.06 to 0.05 | 0.47 | 0.52 | -0.09 to 0.01 | 0.57 | 0.58 | -0.06 to 0.07 | 0.50 | 0.48 | -0.04 to 0.06 | 0.51 | 0.51 | -0.05 to 0.05 | 0.35 | 0.35 | -0.02 to 0.02 |
| Cuneal (L) | 0.58 | 0.57 | -0.05 to 0.09 | 0.43 | 0.43 | -0.01 to 0.01 | 0.55 | 0.55 | -0.07 to 0.07 | 0.43 | 0.43 | -0.01 to 0.01 | 0.54 | 0.55 | -0.06 to 0.07 | 0.43 | 0.43 | -0.01 to 0.01 |
| Frontorbital (L) | 0.64 | 0.66 | -0.08 to 0.04 | 0.54 | 0.58 | -0.08 to 0.01 | 0.58 | 0.57 | -0.04 to 0.07 | 0.44 | 0.39 | 0.01 to 0.10 | 0.51 | 0.49 | -0.04 to 0.07 | 0.37 | 0.37 | -0.02 to 0.02 |
| aPH (L) | 0.69 | 0.68 | -0.06 to 0.09 | 0.45 | 0.45 | -0.01 to 0.01 | 0.61 | 0.60 | -0.07 to 0.10 | 0.45 | 0.45 | -0.01 to 0.01 | 0.54 | 0.57 | -0.11 to 0.06 | 0.45 | 0.45 | -0.01 to 0.01 |
| pPH (L) | 0.62 | 0.61 | -0.06 to 0.09 | 0.45 | 0.45 | -0.01 to 0.01 | 0.54 | 0.54 | -0.08 to 0.08 | 0.45 | 0.45 | -0.01 to 0.01 | 0.51 | 0.52 | -0.08 to 0.06 | 0.45 | 0.45 | -0.01 to 0.01 |
| Lingual (L) | 0.57 | 0.59 | -0.08 to 0.04 | 0.39 | 0.39 | -0.02 to 0.02 | 0.52 | 0.53 | -0.08 to 0.05 | 0.39 | 0.39 | -0.02 to 0.02 | 0.55 | 0.53 | -0.04 to 0.08 | 0.39 | 0.39 | -0.02 to 0.02 |
| atFusiform (L) | 0.64 | 0.66 | -0.11 to 0.09 | 0.48 | 0.48 | -0.01 to 0.01 | 0.53 | 0.54 | -0.11 to 0.10 | 0.48 | 0.48 | -0.01 to 0.01 | 0.49 | 0.51 | -0.12 to 0.11 | 0.48 | 0.48 | -0.01 to 0.01 |
| ptFusiform (L) | 0.63 | 0.63 | -0.07 to 0.07 | 0.43 | 0.43 | -0.01 to 0.01 | 0.54 | 0.57 | -0.08 to 0.05 | 0.43 | 0.43 | -0.01 to 0.01 | 0.56 | 0.55 | -0.06 to 0.06 | 0.43 | 0.43 | -0.01 to 0.01 |
| toFusiform (L) | 0.62 | 0.63 | -0.07 to 0.07 | 0.42 | 0.41 | -0.01 to 0.04 | 0.54 | 0.56 | -0.08 to 0.05 | 0.41 | 0.41 | -0.01 to 0.02 | 0.57 | 0.56 | -0.04 to 0.07 | 0.41 | 0.41 | -0.01 to 0.02 |
| occFusiform (L) | 0.60 | 0.63 | -0.08 to 0.03 | 0.41 | 0.43 | -0.05 to 0.02 | 0.54 | 0.55 | -0.06 to 0.05 | 0.39 | 0.39 | -0.02 to 0.02 | 0.57 | 0.57 | -0.05 to 0.06 | 0.39 | 0.39 | -0.02 to 0.02 |
| fOperculum (L) | 0.60 | 0.61 | -0.07 to 0.05 | 0.49 | 0.44 | 0.01 to 0.09 | 0.54 | 0.53 | -0.06 to 0.07 | 0.41 | 0.41 | -0.02 to 0.02 | 0.54 | 0.52 | -0.03 to 0.09 | 0.41 | 0.41 | -0.02 to 0.02 |
| cOperculum (L) | 0.56 | 0.57 | -0.07 to 0.07 | 0.43 | 0.42 | -0.02 to 0.04 | 0.53 | 0.52 | -0.05 to 0.07 | 0.40 | 0.40 | -0.02 to 0.02 | 0.53 | 0.52 | -0.05 to 0.06 | 0.40 | 0.40 | -0.02 to 0.02 |
| pOperculum (L) | 0.59 | 0.59 | -0.06 to 0.07 | 0.41 | 0.42 | -0.03 to 0.01 | 0.55 | 0.55 | -0.06 to 0.08 | 0.41 | 0.41 | -0.02 to 0.02 | 0.56 | 0.54 | -0.04 to 0.10 | 0.41 | 0.41 | -0.02 to 0.02 |
| PP (L) | 0.58 | 0.59 | -0.08 to 0.07 | 0.44 | 0.44 | -0.01 to 0.01 | 0.53 | 0.50 | -0.05 to 0.10 | 0.44 | 0.44 | -0.01 to 0.01 | 0.53 | 0.50 | -0.03 to 0.09 | 0.44 | 0.44 | -0.01 to 0.01 |
| Heschl (L) | 0.60 | 0.61 | -0.07 to 0.09 | 0.44 | 0.49 | -0.08 to -0.01 | 0.53 | 0.52 | -0.06 to 0.08 | 0.44 | 0.44 | -0.01 to 0.02 | 0.57 | 0.55 | -0.03 to 0.11 | 0.44 | 0.44 | -0.01 to 0.02 |
| PT (L) | 0.61 | 0.62 | -0.06 to 0.06 | 0.57 | 0.57 | -0.06 to 0.06 | 0.55 | 0.54 | -0.05 to 0.08 | 0.41 | 0.41 | -0.02 to 0.02 | 0.56 | 0.55 | -0.05 to 0.07 | 0.41 | 0.41 | -0.02 to 0.02 |
| Supracalcarine (L) | 0.58 | 0.58 | -0.08 to 0.07 | 0.44 | 0.44 | -0.01 to 0.01 | 0.53 | 0.53 | -0.08 to 0.07 | 0.44 | 0.44 | -0.01 to 0.01 | 0.51 | 0.51 | -0.06 to 0.07 | 0.44 | 0.44 | -0.01 to 0.01 |
| OP (L) | 0.59 | 0.62 | -0.09 to 0.05 | 0.41 | 0.41 | -0.02 to 0.02 | 0.55 | 0.56 | -0.07 to 0.06 | 0.41 | 0.41 | -0.02 to 0.02 | 0.58 | 0.57 | -0.04 to 0.09 | 0.41 | 0.41 | -0.02 to 0.02 |
| Thalamus (L) | 0.53 | 0.54 | -0.07 to 0.07 | 0.42 | 0.42 | -0.02 to 0.02 | 0.53 | 0.53 | -0.06 to 0.06 | 0.42 | 0.42 | -0.02 to 0.02 | 0.51 | 0.47 | -0.02 to 0.10 | 0.42 | 0.42 | -0.02 to 0.02 |
| Caudate (L) | 0.61 | 0.64 | -0.10 to 0.05 | 0.45 | 0.45 | -0.01 to 0.01 | 0.60 | 0.61 | -0.09 to 0.07 | 0.45 | 0.45 | -0.01 to 0.01 | 0.53 | 0.52 | -0.08 to 0.09 | 0.45 | 0.45 | -0.01 to 0.01 |
| Putamen (L) | 0.59 | 0.59 | -0.07 to 0.06 | 0.44 | 0.44 | -0.01 to 0.02 | 0.57 | 0.58 | -0.08 to 0.05 | 0.44 | 0.44 | -0.01 to 0.02 | 0.49 | 0.48 | -0.06 to 0.08 | 0.44 | 0.44 | -0.01 to 0.02 |
| Pallidum (L) | 0.59 | 0.58 | -0.07 to 0.10 | 0.46 | 0.46 | -0.01 to 0.01 | 0.56 | 0.57 | -0.09 to 0.07 | 0.46 | 0.46 | -0.01 to 0.01 | 0.51 | 0.48 | -0.04 to 0.12 | 0.46 | 0.46 | -0.01 to 0.01 |
| Hippocampus (L) | 0.67 | 0.65 | -0.05 to 0.08 | 0.44 | 0.44 | -0.01 to 0.01 | 0.60 | 0.58 | -0.06 to 0.08 | 0.44 | 0.44 | -0.01 to 0.01 | 0.53 | 0.54 | -0.07 to 0.07 | 0.44 | 0.44 | -0.01 to 0.01 |
| Amygdala (L) | 0.69 | 0.68 | -0.07 to 0.09 | 0.45 | 0.45 | -0.01 to 0.01 | 0.64 | 0.64 | -0.07 to 0.09 | 0.45 | 0.45 | -0.01 to 0.01 | 0.54 | 0.56 | -0.10 to 0.07 | 0.45 | 0.45 | -0.01 to 0.01 |
| Accumbens (L) | 0.66 | 0.67 | -0.10 to 0.10 | 0.48 | 0.48 | -0.01 to 0.01 | 0.62 | 0.65 | -0.14 to 0.09 | 0.48 | 0.48 | -0.01 to 0.01 | 0.48 | 0.52 | -0.13 to 0.10 | 0.48 | 0.48 | -0.01 to 0.01 |
| Brainstem (L) | 0.57 | 0.57 | -0.09 to 0.13 | 0.47 | 0.47 | -0.01 to 0.01 | 0.55 | 0.54 | -0.06 to 0.11 | 0.47 | 0.47 | -0.01 to 0.01 | 0.50 | 0.50 | -0.09 to 0.09 | 0.47 | 0.47 | -0.01 to 0.01 |
| Cerebellum (L) | 0.55 | 0.55 | -0.06 to 0.06 | 0.40 | 0.40 | -0.02 to 0.02 | 0.55 | 0.55 | -0.06 to 0.06 | 0.40 | 0.40 | -0.02 to 0.02 | 0.53 | 0.54 | -0.06 to 0.05 | 0.40 | 0.40 | -0.02 to 0.02 |

**Extended Data Fig. 17 | Left hemispheric forward inference results across classification architectures.** Observed test set ROC-AUC and F1 scores for logistic regression (LR) and neural network (NN) classifiers. Each 99.9% confidence interval (CI) for the difference between LR and NN classifiers was computed by bootstrap resampling of articles in the test set over 1,000 iterations. Classifiers for which the 99.9% CI did not include zero are highlighted in yellow.

| | Data-Driven | | | | | | RDoC | | | | | | DSM | | | | | |
|---|---|---|---|---|---|---|---|---|---|---|---|---|---|---|---|---|---|---|
| | ROC-AUC | | | F1 | | | ROC-AUC | | | F1 | | | ROC-AUC | | | F1 | | |
| | LR | NN | 99.9% CI | LR | NN | 99.9% CI | LR | NN | 99.9% CI | LR | NN | 99.9% CI | LR | NN | 99.9% CI | LR | NN | 99.9% CI |
| FP (R) | 0.62 | 0.62 | -0.06 to 0.06 | 0.59 | 0.59 | -0.05 to 0.06 | 0.60 | 0.60 | -0.06 to 0.07 | 0.57 | 0.57 | -0.04 to 0.06 | 0.54 | 0.54 | -0.06 to 0.05 | 0.35 | 0.35 | -0.02 to 0.02 |
| Insula (R) | 0.61 | 0.61 | -0.05 to 0.06 | 0.40 | 0.42 | -0.05 to 0.02 | 0.59 | 0.59 | -0.05 to 0.06 | 0.41 | 0.38 | 0.01 to 0.06 | 0.51 | 0.51 | -0.05 to 0.06 | 0.38 | 0.38 | -0.02 to 0.02 |
| SFG (R) | 0.59 | 0.60 | -0.06 to 0.05 | 0.57 | 0.57 | -0.05 to 0.07 | 0.60 | 0.61 | -0.06 to 0.05 | 0.58 | 0.58 | -0.05 to 0.06 | 0.52 | 0.51 | -0.04 to 0.07 | 0.50 | 0.49 | -0.05 to 0.07 |
| MFG (R) | 0.61 | 0.61 | -0.06 to 0.05 | 0.59 | 0.59 | -0.04 to 0.06 | 0.62 | 0.62 | -0.05 to 0.06 | 0.59 | 0.59 | -0.04 to 0.05 | 0.55 | 0.53 | -0.04 to 0.07 | 0.52 | 0.50 | -0.05 to 0.07 |
| trIFG (R) | 0.63 | 0.64 | -0.06 to 0.05 | 0.41 | 0.46 | -0.09 to 0.00 | 0.58 | 0.58 | -0.06 to 0.07 | 0.42 | 0.39 | 0.00 to 0.06 | 0.54 | 0.52 | -0.05 to 0.07 | 0.39 | 0.39 | -0.02 to 0.02 |
| opIFG (R) | 0.60 | 0.60 | -0.06 to 0.05 | 0.50 | 0.53 | -0.07 to 0.03 | 0.57 | 0.57 | -0.05 to 0.07 | 0.51 | 0.50 | -0.05 to 0.06 | 0.55 | 0.55 | -0.05 to 0.05 | 0.37 | 0.37 | -0.02 to 0.02 |
| Precentral (R) | 0.58 | 0.60 | -0.07 to 0.04 | 0.57 | 0.57 | -0.05 to 0.05 | 0.61 | 0.62 | -0.07 to 0.06 | 0.58 | 0.58 | -0.04 to 0.06 | 0.53 | 0.55 | -0.07 to 0.05 | 0.48 | 0.45 | -0.02 to 0.07 |
| TP (R) | 0.60 | 0.60 | -0.06 to 0.08 | 0.43 | 0.43 | -0.01 to 0.02 | 0.58 | 0.58 | -0.07 to 0.06 | 0.43 | 0.43 | -0.01 to 0.02 | 0.52 | 0.53 | -0.08 to 0.05 | 0.43 | 0.43 | -0.01 to 0.02 |
| aSTG (R) | 0.61 | 0.61 | -0.08 to 0.09 | 0.45 | 0.45 | -0.01 to 0.01 | 0.57 | 0.56 | -0.06 to 0.09 | 0.45 | 0.45 | -0.01 to 0.01 | 0.52 | 0.50 | -0.06 to 0.08 | 0.45 | 0.45 | -0.01 to 0.01 |
| pSTG (R) | 0.60 | 0.60 | -0.06 to 0.07 | 0.51 | 0.54 | -0.08 to 0.03 | 0.57 | 0.56 | -0.05 to 0.09 | 0.41 | 0.41 | -0.02 to 0.02 | 0.53 | 0.53 | -0.06 to 0.06 | 0.41 | 0.41 | -0.02 to 0.02 |
| aMTG (R) | 0.61 | 0.61 | -0.08 to 0.10 | 0.46 | 0.46 | -0.01 to 0.01 | 0.61 | 0.61 | -0.07 to 0.07 | 0.46 | 0.46 | -0.01 to 0.01 | 0.50 | 0.49 | -0.07 to 0.08 | 0.46 | 0.46 | -0.01 to 0.01 |
| pMTG (R) | 0.61 | 0.61 | -0.07 to 0.07 | 0.46 | 0.44 | -0.02 to 0.05 | 0.58 | 0.58 | -0.07 to 0.06 | 0.42 | 0.42 | -0.01 to 0.02 | 0.52 | 0.52 | -0.06 to 0.06 | 0.42 | 0.42 | -0.01 to 0.02 |
| toMTG (R) | 0.59 | 0.59 | -0.05 to 0.07 | 0.41 | 0.40 | -0.01 to 0.05 | 0.58 | 0.57 | -0.05 to 0.07 | 0.39 | 0.39 | -0.02 to 0.02 | 0.54 | 0.54 | -0.06 to 0.06 | 0.39 | 0.39 | -0.02 to 0.02 |
| aITG (R) | 0.58 | 0.61 | -0.13 to 0.10 | 0.48 | 0.48 | -0.01 to 0.01 | 0.57 | 0.59 | -0.13 to 0.09 | 0.48 | 0.48 | -0.01 to 0.01 | 0.48 | 0.51 | -0.13 to 0.07 | 0.48 | 0.48 | -0.01 to 0.01 |
| pITG (R) | 0.58 | 0.58 | -0.08 to 0.08 | 0.46 | 0.46 | -0.01 to 0.01 | 0.57 | 0.58 | -0.08 to 0.09 | 0.46 | 0.46 | -0.01 to 0.01 | 0.51 | 0.54 | -0.11 to 0.05 | 0.46 | 0.46 | -0.01 to 0.01 |
| toITG (R) | 0.62 | 0.62 | -0.07 to 0.06 | 0.43 | 0.43 | -0.02 to 0.02 | 0.57 | 0.56 | -0.05 to 0.07 | 0.43 | 0.43 | -0.02 to 0.02 | 0.55 | 0.56 | -0.06 to 0.05 | 0.43 | 0.43 | -0.02 to 0.02 |
| Postcentral (R) | 0.56 | 0.57 | -0.07 to 0.04 | 0.54 | 0.54 | -0.05 to 0.05 | 0.57 | 0.57 | -0.06 to 0.07 | 0.55 | 0.55 | -0.06 to 0.05 | 0.52 | 0.52 | -0.06 to 0.05 | 0.36 | 0.36 | -0.02 to 0.02 |
| SPL (R) | 0.61 | 0.62 | -0.07 to 0.05 | 0.50 | 0.51 | -0.06 to 0.05 | 0.60 | 0.60 | -0.05 to 0.07 | 0.50 | 0.50 | -0.04 to 0.06 | 0.55 | 0.54 | -0.04 to 0.06 | 0.38 | 0.38 | -0.02 to 0.02 |
| aSMG (R) | 0.57 | 0.58 | -0.07 to 0.05 | 0.41 | 0.41 | -0.02 to 0.02 | 0.59 | 0.59 | -0.06 to 0.07 | 0.41 | 0.41 | -0.02 to 0.02 | 0.54 | 0.53 | -0.05 to 0.06 | 0.41 | 0.41 | -0.02 to 0.02 |
| pSMG (R) | 0.59 | 0.60 | -0.08 to 0.05 | 0.58 | 0.57 | -0.04 to 0.06 | 0.59 | 0.59 | -0.06 to 0.05 | 0.57 | 0.57 | -0.04 to 0.05 | 0.56 | 0.56 | -0.05 to 0.06 | 0.54 | 0.54 | -0.05 to 0.05 |
| Angular (R) | 0.60 | 0.62 | -0.07 to 0.04 | 0.58 | 0.58 | -0.05 to 0.05 | 0.61 | 0.60 | -0.06 to 0.06 | 0.57 | 0.59 | -0.07 to 0.04 | 0.56 | 0.56 | -0.05 to 0.06 | 0.53 | 0.53 | -0.05 to 0.05 |
| slOccipital (R) | 0.63 | 0.65 | -0.08 to 0.04 | 0.60 | 0.61 | -0.05 to 0.05 | 0.60 | 0.60 | -0.06 to 0.06 | 0.57 | 0.58 | -0.06 to 0.04 | 0.55 | 0.55 | -0.05 to 0.06 | 0.53 | 0.52 | -0.05 to 0.06 |
| ilOccipital (R) | 0.62 | 0.63 | -0.06 to 0.05 | 0.54 | 0.58 | -0.09 to 0.01 | 0.57 | 0.58 | -0.06 to 0.06 | 0.41 | 0.43 | -0.06 to 0.02 | 0.56 | 0.55 | -0.04 to 0.06 | 0.37 | 0.36 | -0.02 to 0.02 |
| Intracalcarine (R) | 0.57 | 0.58 | -0.07 to 0.05 | 0.43 | 0.43 | -0.01 to 0.01 | 0.54 | 0.53 | -0.05 to 0.10 | 0.43 | 0.43 | -0.01 to 0.01 | 0.54 | 0.53 | -0.05 to 0.08 | 0.43 | 0.43 | -0.01 to 0.01 |
| Frontomedial (R) | 0.61 | 0.61 | -0.07 to 0.07 | 0.46 | 0.46 | -0.01 to 0.01 | 0.62 | 0.62 | -0.08 to 0.09 | 0.46 | 0.46 | -0.01 to 0.01 | 0.55 | 0.58 | -0.10 to 0.07 | 0.46 | 0.46 | -0.01 to 0.01 |
| SMA (R) | 0.56 | 0.58 | -0.08 to 0.04 | 0.41 | 0.41 | -0.01 to 0.02 | 0.58 | 0.59 | -0.07 to 0.07 | 0.41 | 0.41 | -0.01 to 0.02 | 0.54 | 0.54 | -0.05 to 0.06 | 0.41 | 0.41 | -0.01 to 0.02 |
| Subcallosal (R) | 0.63 | 0.61 | -0.07 to 0.14 | 0.47 | 0.47 | -0.01 to 0.01 | 0.63 | 0.63 | -0.09 to 0.14 | 0.47 | 0.47 | -0.01 to 0.01 | 0.54 | 0.56 | -0.11 to 0.08 | 0.47 | 0.47 | -0.01 to 0.01 |
| Paracingulate (R) | 0.60 | 0.60 | -0.05 to 0.06 | 0.57 | 0.58 | -0.05 to 0.05 | 0.59 | 0.59 | -0.04 to 0.06 | 0.55 | 0.55 | -0.05 to 0.05 | 0.49 | 0.52 | -0.07 to 0.03 | 0.37 | 0.38 | -0.05 to 0.02 |
| ACC (R) | 0.60 | 0.60 | -0.06 to 0.06 | 0.58 | 0.58 | -0.05 to 0.05 | 0.60 | 0.60 | -0.05 to 0.07 | 0.52 | 0.53 | -0.05 to 0.05 | 0.49 | 0.50 | -0.06 to 0.04 | 0.36 | 0.38 | -0.05 to 0.01 |
| PCC (R) | 0.56 | 0.57 | -0.06 to 0.05 | 0.40 | 0.40 | -0.02 to 0.02 | 0.57 | 0.56 | -0.07 to 0.06 | 0.40 | 0.40 | -0.02 to 0.02 | 0.51 | 0.51 | -0.05 to 0.06 | 0.40 | 0.40 | -0.02 to 0.02 |
| Precuneous (R) | 0.57 | 0.56 | -0.05 to 0.08 | 0.47 | 0.51 | -0.09 to 0.01 | 0.57 | 0.58 | -0.06 to 0.06 | 0.50 | 0.51 | -0.05 to 0.05 | 0.53 | 0.52 | -0.05 to 0.05 | 0.36 | 0.36 | -0.02 to 0.02 |
| Cuneal (R) | 0.59 | 0.58 | -0.05 to 0.06 | 0.42 | 0.42 | -0.02 to 0.02 | 0.56 | 0.55 | -0.06 to 0.08 | 0.42 | 0.42 | -0.02 to 0.02 | 0.54 | 0.53 | -0.04 to 0.07 | 0.42 | 0.42 | -0.02 to 0.02 |
| Frontorbital (R) | 0.62 | 0.63 | -0.06 to 0.05 | 0.55 | 0.56 | -0.06 to 0.04 | 0.59 | 0.59 | -0.05 to 0.06 | 0.49 | 0.46 | -0.01 to 0.08 | 0.52 | 0.50 | -0.04 to 0.06 | 0.37 | 0.37 | -0.02 to 0.02 |
| aPH (R) | 0.66 | 0.65 | -0.06 to 0.09 | 0.46 | 0.46 | -0.01 to 0.01 | 0.61 | 0.61 | -0.07 to 0.09 | 0.46 | 0.46 | -0.01 to 0.01 | 0.55 | 0.57 | -0.10 to 0.07 | 0.46 | 0.46 | -0.01 to 0.01 |
| pPH (R) | 0.65 | 0.63 | -0.08 to 0.10 | 0.46 | 0.46 | -0.01 to 0.01 | 0.54 | 0.55 | -0.08 to 0.08 | 0.46 | 0.46 | -0.01 to 0.01 | 0.50 | 0.47 | -0.05 to 0.13 | 0.46 | 0.46 | -0.01 to 0.01 |
| Lingual (R) | 0.58 | 0.59 | -0.08 to 0.05 | 0.41 | 0.39 | -0.01 to 0.04 | 0.54 | 0.54 | -0.07 to 0.05 | 0.39 | 0.39 | -0.02 to 0.02 | 0.53 | 0.51 | -0.04 to 0.07 | 0.39 | 0.39 | -0.02 to 0.02 |
| atFusiform (R) | 0.60 | 0.62 | -0.13 to 0.09 | 0.48 | 0.48 | -0.01 to 0.01 | 0.56 | 0.55 | -0.11 to 0.13 | 0.48 | 0.48 | -0.01 to 0.01 | 0.53 | 0.52 | -0.11 to 0.13 | 0.48 | 0.48 | -0.01 to 0.01 |
| ptFusiform (R) | 0.62 | 0.63 | -0.07 to 0.08 | 0.45 | 0.45 | -0.01 to 0.01 | 0.56 | 0.57 | -0.09 to 0.06 | 0.45 | 0.45 | -0.01 to 0.01 | 0.54 | 0.52 | -0.04 to 0.10 | 0.45 | 0.45 | -0.01 to 0.01 |
| toFusiform (R) | 0.64 | 0.63 | -0.05 to 0.07 | 0.42 | 0.42 | -0.02 to 0.02 | 0.55 | 0.56 | -0.07 to 0.07 | 0.42 | 0.42 | -0.02 to 0.02 | 0.54 | 0.55 | -0.06 to 0.05 | 0.42 | 0.42 | -0.02 to 0.02 |
| occFusiform (R) | 0.60 | 0.61 | -0.06 to 0.06 | 0.40 | 0.40 | -0.02 to 0.02 | 0.55 | 0.55 | -0.06 to 0.06 | 0.40 | 0.40 | -0.02 to 0.02 | 0.57 | 0.55 | -0.04 to 0.07 | 0.40 | 0.40 | -0.02 to 0.02 |
| fOperculum (R) | 0.59 | 0.59 | -0.06 to 0.07 | 0.40 | 0.40 | -0.01 to 0.01 | 0.58 | 0.57 | -0.05 to 0.07 | 0.40 | 0.40 | -0.01 to 0.01 | 0.52 | 0.50 | -0.05 to 0.07 | 0.40 | 0.40 | -0.01 to 0.01 |
| cOperculum (R) | 0.56 | 0.57 | -0.07 to 0.06 | 0.41 | 0.41 | -0.02 to 0.02 | 0.56 | 0.54 | -0.04 to 0.09 | 0.41 | 0.41 | -0.02 to 0.02 | 0.54 | 0.53 | -0.04 to 0.07 | 0.41 | 0.41 | -0.02 to 0.02 |
| pOperculum (R) | 0.54 | 0.54 | -0.08 to 0.07 | 0.43 | 0.43 | -0.01 to 0.01 | 0.55 | 0.53 | -0.06 to 0.09 | 0.43 | 0.43 | -0.01 to 0.01 | 0.55 | 0.54 | -0.04 to 0.07 | 0.43 | 0.43 | -0.01 to 0.01 |
| PP (R) | 0.58 | 0.57 | -0.06 to 0.09 | 0.44 | 0.44 | -0.01 to 0.01 | 0.53 | 0.49 | -0.03 to 0.12 | 0.44 | 0.44 | -0.01 to 0.01 | 0.53 | 0.49 | -0.03 to 0.10 | 0.44 | 0.44 | -0.01 to 0.01 |
| Heschl (R) | 0.57 | 0.57 | -0.06 to 0.09 | 0.45 | 0.45 | -0.01 to 0.01 | 0.54 | 0.51 | -0.05 to 0.10 | 0.45 | 0.45 | -0.01 to 0.01 | 0.55 | 0.52 | -0.03 to 0.09 | 0.45 | 0.45 | -0.01 to 0.01 |
| PT (R) | 0.58 | 0.59 | -0.07 to 0.06 | 0.42 | 0.47 | -0.08 to -0.02 | 0.57 | 0.55 | -0.04 to 0.09 | 0.42 | 0.42 | -0.01 to 0.01 | 0.54 | 0.54 | -0.05 to 0.06 | 0.42 | 0.42 | -0.01 to 0.01 |
| Supracalcarine (R) | 0.57 | 0.57 | -0.06 to 0.06 | 0.43 | 0.43 | -0.01 to 0.01 | 0.55 | 0.54 | -0.06 to 0.10 | 0.43 | 0.43 | -0.01 to 0.01 | 0.54 | 0.53 | -0.05 to 0.07 | 0.43 | 0.43 | -0.01 to 0.01 |
| OP (R) | 0.60 | 0.62 | -0.08 to 0.05 | 0.41 | 0.41 | -0.01 to 0.02 | 0.54 | 0.55 | -0.08 to 0.06 | 0.41 | 0.41 | -0.01 to 0.02 | 0.55 | 0.54 | -0.04 to 0.06 | 0.41 | 0.41 | -0.01 to 0.02 |
| Thalamus (R) | 0.54 | 0.53 | -0.06 to 0.07 | 0.42 | 0.42 | -0.02 to 0.02 | 0.56 | 0.56 | -0.08 to 0.07 | 0.42 | 0.42 | -0.02 to 0.02 | 0.53 | 0.48 | -0.01 to 0.12 | 0.42 | 0.42 | -0.02 to 0.02 |
| Caudate (R) | 0.58 | 0.59 | -0.09 to 0.06 | 0.45 | 0.45 | -0.01 to 0.01 | 0.57 | 0.59 | -0.10 to 0.06 | 0.45 | 0.45 | -0.01 to 0.01 | 0.50 | 0.52 | -0.10 to 0.06 | 0.45 | 0.45 | -0.01 to 0.01 |
| Putamen (R) | 0.60 | 0.61 | -0.08 to 0.06 | 0.44 | 0.44 | -0.01 to 0.01 | 0.61 | 0.60 | -0.06 to 0.08 | 0.44 | 0.44 | -0.01 to 0.01 | 0.56 | 0.52 | -0.02 to 0.11 | 0.44 | 0.44 | -0.01 to 0.01 |
| Pallidum (R) | 0.58 | 0.57 | -0.08 to 0.10 | 0.46 | 0.46 | -0.01 to 0.01 | 0.57 | 0.58 | -0.09 to 0.07 | 0.46 | 0.46 | -0.01 to 0.01 | 0.52 | 0.49 | -0.05 to 0.11 | 0.46 | 0.46 | -0.01 to 0.01 |
| Hippocampus (R) | 0.66 | 0.65 | -0.06 to 0.07 | 0.44 | 0.44 | -0.01 to 0.02 | 0.59 | 0.60 | -0.06 to 0.06 | 0.44 | 0.44 | -0.01 to 0.02 | 0.53 | 0.54 | -0.08 to 0.08 | 0.44 | 0.44 | -0.01 to 0.02 |
| Amygdala (R) | 0.69 | 0.67 | -0.04 to 0.11 | 0.45 | 0.45 | -0.01 to 0.01 | 0.66 | 0.65 | -0.07 to 0.10 | 0.45 | 0.45 | -0.01 to 0.01 | 0.57 | 0.59 | -0.09 to 0.07 | 0.45 | 0.45 | -0.01 to 0.01 |
| Accumbens (R) | 0.64 | 0.64 | -0.12 to 0.10 | 0.48 | 0.48 | -0.01 to 0.01 | 0.64 | 0.64 | -0.13 to 0.14 | 0.48 | 0.48 | -0.01 to 0.01 | 0.48 | 0.50 | -0.13 to 0.08 | 0.48 | 0.48 | -0.01 to 0.01 |
| Brainstem (R) | 0.56 | 0.57 | -0.10 to 0.08 | 0.47 | 0.47 | -0.01 to 0.01 | 0.55 | 0.55 | -0.09 to 0.10 | 0.47 | 0.47 | -0.01 to 0.01 | 0.50 | 0.46 | -0.06 to 0.13 | 0.47 | 0.47 | -0.01 to 0.01 |
| Cerebellum (R) | 0.55 | 0.55 | -0.05 to 0.07 | 0.40 | 0.40 | -0.02 to 0.02 | 0.56 | 0.56 | -0.05 to 0.08 | 0.40 | 0.40 | -0.02 to 0.02 | 0.54 | 0.52 | -0.04 to 0.08 | 0.40 | 0.40 | -0.02 to 0.02 |

**Extended Data Fig. 18 | Right hemispheric forward inference results across classification architectures.** Observed test set ROC-AUC and F1 scores for logistic regression (LR) and neural network (NN) classifiers. Each 99.9% confidence interval (CI) for the difference between LR and NN classifiers was computed by bootstrap resampling of articles in the test set over 1,000 iterations. In all cases, the 99.9% CI included zero.

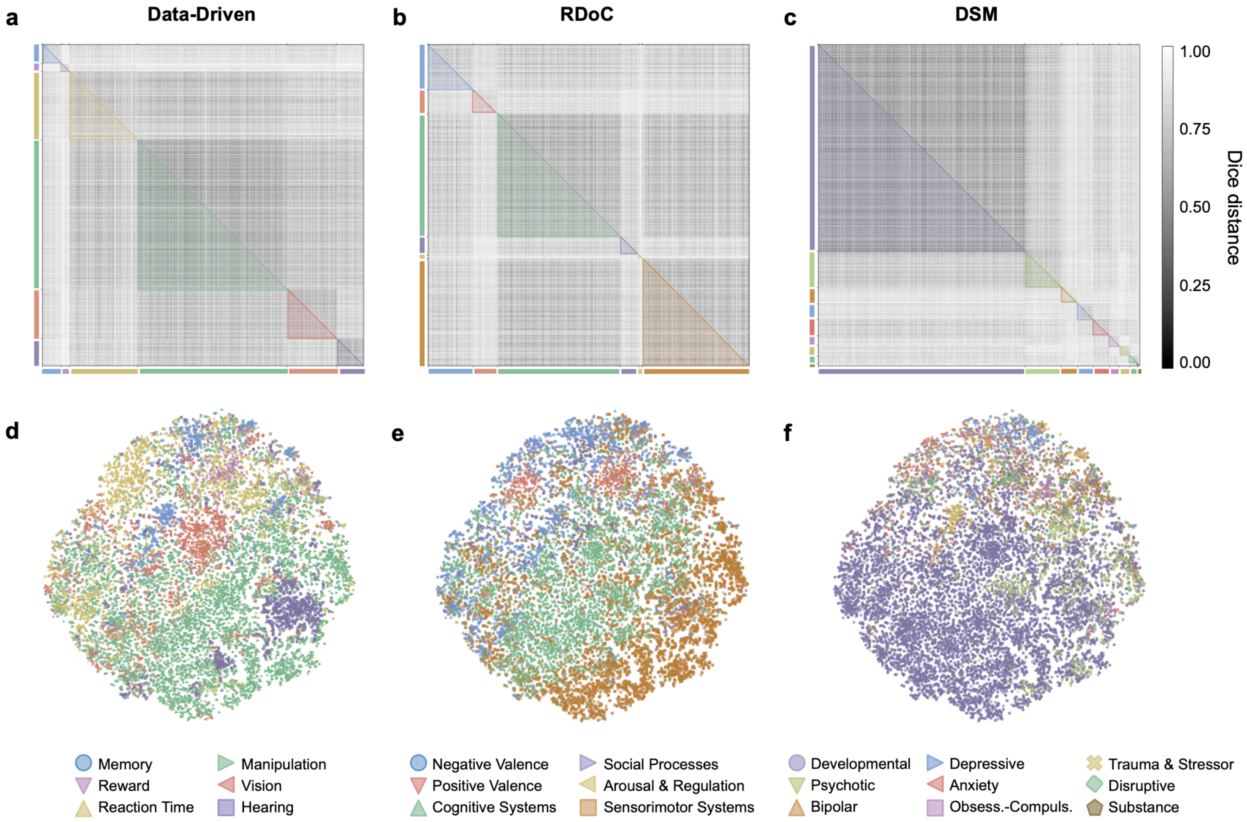


**Extended Data Fig. 19 | Articles partitioned by similarity to mental functions and brain circuits in the domains of each framework.** Dice distance between articles (*n* = 18,155) is shown for binarized vectors of the mental function words that occurred in the full text and the brain structures to which reported coordinate data were mapped. Articles were matched to domains based on the dice similarity of their word-structure vectors, and domain assignments are represented by the color coding scheme established in Fig. 3 for **a,** the data-driven ontology, **b,** RDoC, and **c,** the DSM. Shaded areas represent the lower triangle of distances between articles within each domain partition. **d-f,** Dice distance between articles across the corpus (*n* = 18,155) visualized with t-SNE. Distances were computed between the terms and structures of every article, and dimensionality of the 18,155 x 18,155 matrix was reduced by principal component analysis. The first 10 principal components (18,155 x 10) were taken as inputs to t-SNE, which was trained with perplexity = 25, early exaggeration = 15, learning rate = 500, and maximum iterations = 1,000. Articles are colored by their domain assignments in each framework.
